# CXXC-finger protein 1 associates with FOXP3 to stabilize homeostasis and suppressive functions of regulatory T cells

**DOI:** 10.1101/2024.09.30.615805

**Authors:** Xiaoyu Meng, Yezhang Zhu, Kuai Liu, Yuxi Wang, Xiaoqian Liu, Chenxin Liu, Yan Zeng, Shuai Wang, Xianzhi Gao, Xin Shen, Jing Chen, Sijue Tao, Qianying Xu, Linjia Dong, Li Shen, Lie Wang

## Abstract

FOXP3-expressing regulatory T (T_reg_) cells play a pivotal role in maintaining immune homeostasis and tolerance, with their activation being crucial for preventing various inflammatory responses. However, the mechanisms governing the epigenetic program in T_reg_ cells during their dynamic activation remain unclear. In this study, we demonstrate that CXXC finger protein 1 (CXXC1) interacts with the transcription factor FOXP3 and facilitates the regulation of target genes by modulating H3K4me3 deposition. *Cxxc1* deletion in T_reg_ cells leads to severe inflammatory disease and spontaneous T-cell activation, with impaired immunosuppressive function. As a transcriptional regulator, CXXC1 promotes the expression of key T_reg_ functional markers under steady-state conditions, which are essential for the maintenance of T_reg_ cell homeostasis and their suppressive functions. Epigenetically, CXXC1 binds to the genomic regulatory regions of T_reg_ program genes in mouse T_reg_ cells, overlapping with FOXP3 binding sites. Given its critical role in T_reg_ cell homeostasis, CXXC1 presents itself as a promising therapeutic target for autoimmune diseases.

## Introduction

Regulatory T (T_reg_) cells are a distinct subset of CD4^+^ T cells that play a critical role in maintaining immune homeostasis and self-tolerance by suppressing excessive or aberrant immune responses to foreign or self-antigens ^1–3^. These cells can be further categorized into thymus-derived regulatory T cells (tT_reg_ cells), periphery-derived T_reg_ cells (pT_reg_ cells), and induced T_reg_ cells (iT_reg_ cells) ^4^. They uniquely express the transcription factor FOXP3, a member of the forkhead winged-helix family, which is essential for T_reg_ cell lineage commitment and suppressive function ^5,6^. Deletion or mutation of the *Foxp3* gene leads to a range of immunological disorders, including allergies, immunopathology, and autoimmune diseases in both mice and humans ^7,8^.

It is well established that FOXP3 recruits various cofactors to form complexes that either promote or repress the expression of downstream genes, with histone and DNA modifications playing pivotal roles in this process. FOXP3 can activate or repress the transcription of key regulators of T_reg_ cell activation and function by recruiting the histone acetyltransferases (HATs) or histone deacetylases (HDACs)^9,10^. Notably, FOXP3-bound sites exhibit enrichment of H3K27me3, a modification essential for FOXP3-mediated repressive chromatin remodeling under inflammatory conditions^11^. However, the direct role of FOXP3 as a transcriptional activator through interactions with epigenetic regulators, particularly via modulation of H3K4 trimethylation, remains poorly documented.

In mammals, six proteins have been identified that catalyze H3K4 methylation. These proteins contain the SET domain and include MLL1 (KMT2A), MLL2 (KMT2B), MLL3 (KMT2C), MLL4 (KMT2D), SETD1A, and SETD1B ^12,13^. For example, MLL1 plays a critical role as an epigenetic regulator in T_reg_ cell activation and functional specialization^14^. Additionally, Placek *et al.* demonstrated that MLL4 is essential for T_reg_ cell development by catalyzing H3K4me1 at distant unbound enhancers through chromatin looping ^15^. H3K4me3, which is enriched at the transcription start site (TSS) and the CpG island (CGI), converts chromatin into active euchromatin by recruiting activating factors ^16^. CXXC finger protein 1(CXXC1, also known as CFP1), which contains a SET1 interaction domain (SID), is required for binding to the histone H3K4 methyltransferases SETD1A and SETD1B ^17^. Previous studies have shown that CXXC1 regulates promoter patterns during T cell maturation, GM-CSF-derived macrophage phagocytosis, TH17 cell differentiation, and the function of ILC3 cells with aging by modulating H3K4me3 modification^18–21^. Despite the well-documented role of CXXC1 in various immune effector cells, its role in T_reg_ cells remains unclear.

Here, we demonstrate that CXXC1 interacts with FOXP3 and enhances the expression of FOXP3 target genes. T_reg_ cell-specific deletion of *Cxxc1* triggers systemic autoimmunity, accompanied by multiorgan inflammation, ultimately resulting in early-onset fatal inflammatory disease in mice. *Cxxc1-*deficient T_reg_ cells exhibit a disadvantage in proliferation and homeostasis, even in non-inflammatory mice where coexisting wild-type (WT) T_reg_ cells were present. Moreover, *Cxxc1*-deficient T_reg_ cells display intrinsic defects in the expression of key suppression molecules, including CTLA-4, CD25, ICOS, and GITR. Consistent with these findings, mechanistic studies strongly suggest that CXXC1 functions as an essential cofactor of FOXP3, playing a crucial role in sustaining H3K4me3 modifications at key T_reg_ cell genes.

## Materials and Methods

### Mice

All mice used in this study were bred for a minimum of seven generations on a C57BL/6 background. Mouse experiments, or cells from mice of the same genotype, compared littermates or age-matched control animals. The *Cxxc1*^fl/fl^ mouse strain has been previously described ^22^. The *Foxp3*^cre^ mice (JAX,016959) were generously provided by Bin Li (Shanghai Jiao Tong University School of Medicine, Shanghai, China). CD45.1 (NM-KI-210226) mice were purchased from the Nanjing Biomedical Research Institute of Nanjing University. *Rag1^−/-^*mice (stock#T004753) were purchased from GemPharmatech. 2D2 (MOG35-55-specific TCR transgenic) mice were graciously supplied by Prof. Linrong Lu (Zhejiang University School of Medicine, Hangzhou, Zhejiang, China). *Foxp3*^cre^ (WT) and *Foxp3*^cre^ *Cxxc1*^fl/fl^ (cKO) were utilized at 3 weeks old unless otherwise stated. All mice were housed in the Zhejiang University Laboratory Animal Center under specific pathogen-free conditions, and all animal experimental procedures were approved by the Zhejiang University Animal Care and Use Committee (approval no.ZJU20230246).

### Cell culture

HEK 293T cells were obtained from ATCC, and Plat E cells were kindly provided by Prof. Xiaolong Liu (Shanghai Institutes for Biological Sciences). Both cell lines were cultured in Dulbecco-modified Eagle medium (DMEM) containing 10% (v/v) fetal bovine serum (FBS), supplemented with 1% penicillin/streptomycin.

### Immunofluorescence (IF) Staining

As previously described ^23^, coverslips were treated with a 0.01% poly-L-lysine solution (P4707; Sigma) for 10 minutes, air-dried, and then coated with CD4^+^ YFP^+^ T_reg_ cells. The cells were then fixed in 4% formaldehyde for 15 minutes at room temperature, permeabilized with 0.2% Triton X-100, and blocked with 1% BSA. Antibodies against CXXC1 (ab198977; Abcam) and FOXP3( 17-5773-82; Invitrogen) were diluted in Image iT FX signal enhancer (I3693; Invitrogen) and incubated with cells overnight at 4 °C. After washing with phosphate-buffered saline (PBS), the cells were incubated with a goat anti-rabbit antibody Alexa Fluor 594 (1:250;10015289; Invitrogen) secondary antibody and stained with DAPI (200 ng/mL; D523; Dojindo). Slides were washed with PBS and sealed with an antifade solution(P36934; Invitrogen) before imaging with an Olympus FV3000 fluorescence microscope. The images were visualized using the FV31-SW software.

### Co-immunoprecipitation and western blot

Harvest the appropriate transfected cell lines and primary cells from culture and wash them with ice-cold PBS. Lyse the cells in NETN300 buffer (300 mM NaCl, 0.5 mM EDTA, 0.5% (v/v) NP-40, 20 mM Tris-HCl pH 8.0) supplemented with a protease inhibitor (1:100, P8340, Sigma-Aldrich) and PMSF (1 mM) on ice for 10 minutes. Take a small portion of the whole-cell lysate as input, and incubate the remaining lysate with either Anti-FLAG M2 Beads (M8823; Sigma) or Anti-HA Beads (HY-K0201; MCE) on a rotator at 4°C overnight. For the endogenous Co-IP assay targeting CXXC1 and FOXP3, incubate the cell lysate with protein G magnetic beads along with anti-CXXC1 (ab198977; Abcam) or anti-FOXP3 (14-4774-82; Invitrogen) antibodies on a rotator at 4°C overnight. Wash the beads three times with IP buffer (100 mM NaCl, 0.5 mM EDTA, 0.5% (v/v) NP-40, 20 mM Tris-HCl pH 8.0) to remove non-specific binding. Boil the washed beads with 1× Laemmli sample buffer (1610747; Bio-Rad) to elute the bound proteins. Separate the denatured proteins by SDS-PAGE. Transfer the separated proteins onto PVDF membranes (Millipore) for immunoblotting.Immunoblot the PVDF membranes (IPVH00010) with the following antibodies: anti-CXXC1 (1:1000; ab198977; Abcam), anti-FOXP3 (1:500; 14-7979-80; Invitrogen), anti-FlAG (1:1000; 14793; Cell Signaling Technology), anti-HA (1:1000; 3724S; Cell Signaling Technology).Detect the immunoblotted proteins using a secondary HRP-conjugated goat anti-rabbit antibody (1:1000; HA1001-100; Huabio) and visualize the bands using an appropriate detection method.

### ELISA

Serum samples from 3-week-old WT and KO mice were analyzed for total IgG and IgE concentrations using ELISA kits (88-50630-88; eBioscience) according to the manufacturer’s instructions. Half-area ELISA plates were coated with Coating Buffer and incubated overnight at 4°C. After washing with PBST (PBS, 1 mM EDTA, 0.05% Tween-20), the plates were blocked with 5% BSA in PBS for 30 minutes at room temperature. Serum was diluted to the appropriate concentration with blocking buffer and incubated overnight at 4°C. After washing with PBST, the plates were incubated with HRP-conjugated anti-mouse IgG(1033-05; SouthernBiotech) and IgE(1110-05; SouthernBiotech) antibodies (1:2000 in 1% BSA/PBST) at 37°C for 1 hour. Following washing, TMB substrate was added, and the reaction was stopped with 2 M HCSOC after sufficient color development (1–15 minutes). Absorbance at 450 nm was measured within 30 minutes.

### Lymphocyte Isolation and Flow Cytometry

Cells from lymphoid organs were prepared by mechanical disruption between frosted slides, while non-lymphoid organs were processed enzymatically. For lung tissue, minced samples were digested in RPMI containing 100 μg/mL DNase I (9003-98-9;Sigma-Aldrich) and 2 mg/mL Collagenase D (LS004188;Worthington Biochemical) at 37°C for 1.5 hours. Liver tissue was minced and digested in RPMI supplemented with 100 μg/mL DNase I and 1 mg/mL Collagenase D at 37°C for 30 minutes, with lymphocytes isolated using a 40-70% Percoll (GE Healthcare) gradient. For intestinal tissue, the samples were first incubated in DMEM containing 3% FBS, 0.2% HEPES, 0.5 M EDTA, and 0.145 mg/mL dithiothreitol for 10 minutes. This was followed by digestion with 50 mg/mL DNase I and 145 mg/mL Collagenase II (Worthington Biochemical) in DMEM at 37°C for 5 minutes. Lymphocytes were then isolated using an 80% and 40% Percoll gradient.

For surface marker analysis, cells were incubated for 15 minutes with purified anti-mouse CD16/32 antibody (101320; BioLegend) to block Fc receptors. After blocking, cells were stained with the indicated antibodies for surface markers. To determine cytokine expression, cells were stimulated for 4 hours at 37C with phorbol12-myristate13-acetate (50ng/mL; S1819; Beyotime), ionomycin (1mg/ml; S1672; Beyotime), and brefeldin A (BFA;00-4506-51; Invitrogen). After stimulation, cells were labeled with a fixable viability dye and surface markers. Cells were then fixed and permeabilized according to the manufacturer’s instructions (00-8222-49; Invitrogen). For transcription factor staining, samples were fixed using the FOXP3/Transcription Factor Staining Buffer Set (00-5523-00; Invitrogen). Flow cytometry was conducted using a BD Fortessa (BD Biosciences) or LongCyte (Beijing Challen Biotechnology Co., Ltd.) system. Flow cytometry data were acquired and analyzed using FlowJo software.

The following antibodies were purchased from Invitrogen or BioLegend: Zombie Viole t f-ixable viability(423113), Zombie NIR fixable(423105), CD25(PC61), CD8α(53-6.7), CD62L(MEL-14), PD-1(J43), CD44(IM7), IL-17A(TC1118H10), KL-RG1(2F1), CD4 (GK1.5), TCRβ(H57-597), IFN-γ(XMG1.2), FOXP3(FJK-16s), CTLA-4(UC10-4B9), ICOS(15F9), GITR(DTA-1), LAG3(C9B7W), TCRVβ11(RR3-15), CD45RB(C363-16A) C D69 (H1.2F3), MHCII(M5/114.15.2), CD80(16-10A1), T-bet(eBio4B10), IKZF4(ESB7C 2), IL-4(11B11), CCR7(4B12), CD73(TY/11.8).

### Real-time PCR

Total RNA was extracted from T_reg_ cells using the RNAAiso Plus(9109; Takara) reagent according to the manufacturer’s instructions, and cDNA synthesis was performed using the Prime Script RT Reagent Kit (Takara). TB Green Premix Ex Taq (RR420A; Takara) was used for quantitative real-time PCR (qPCR). The expression levels of target mRNA were normalized to the level of β*-actin* expression. The primers for qPCR are as follows:

*Cxxc1* qPCR Forward: ATCCGGGAATGGTACTGTCG
*Cxxc1* qPCR Reverse: CTGTGGAGAAGATTTGTGGG
*β-actin* qPCR Forward: CTGTCCCTGTATGCCTCTG
*β-actin* qPCR Reverse: ATGTCACGCACGATTTCC

### CD4^+^T and YFP^+^ T_reg_ cells adoptive transfer in Experimental autoimmune encephalomyelitis (EAE)

CD4^+^YFP^+^ T_reg_ cells from *Foxp3*^cre^ and *Foxp3*^cre^ *Cxxc1*^fl/fl^ mice were enriched using the Mouse CD4 T Cell Isolation Kit (480005; Biolegend) and then sorted using the BD Aria II flow cytometer. Naïve CD4^+^T cells from 2D2 (MOG35-55-specificTCR transgenic) mice were isolated using the Mouse CD4 Naïve T cell Isolation Kit480039 (480039; Biolegend). As previously described^24^, 2D2 naïve CD4^+^ T cells alone (5 × 10^5^ per mouse), or 2D2 naïve CD4^+^ T cells (5× 10^5^ per mouse) together with WT or *Foxp3*^cre^*Cxxc1*^fl/fl^ T_reg_ cells (2×10^5^ per mouse), were transferred into *Rag1^−/−^* mice via the tail vein. One day after cell transfer, the recipient mice were inoculated subcutaneously (s.c.) with 200μg MOG35-55 peptide (MEVGWYRSPFSRVVHLYRNGK; GenemeSynthesis) emulsified in complete Freund’s adjuvant (CFA) (F5506; Sigma). Intravenous administration of 200 ng of Pertussis toxin (181; List Biological Laboratories) was performed on days 0 and 2 after peptide inoculation. The severity of EAE was monitored and blindly graded using a clinical score from 0 to 5: 0, no clinical signs; 1, limp tail; 2, paraparesis (weakness, incomplete paralysis of one or two hind limbs); 3, paraplegia (complete paralysis of two hind limbs); 4, paraplegia with forelimb weakness or paralysis; 5, dying or death.

### Isolation lymphocytes from the central nervous system (CNS)

On day 14 after EAE induction, mice were perfused with transcardially administered PBS to eliminate contaminating blood cells in the central nervous system (CNS). The forebrain and cerebellum were dissected to expose the spinal cord, which was then carefully removed from the spinal canal. The fresh spinal cord was harvested and cut into 2 mm pieces. The CNS tissue pieces were homogenized using a syringe and passed through a 70 µM cell strainer to obtain a single-cell suspension. The single-cell suspension was digested with collagenase D (2μg/ml;11088858001; Roche) and deoxyribonuclease I (DNase I; 1 μg/ml; DN25; Sigma-Aldrich) at 37°C for 20 minutes under rotation. After digestion, the cell suspension was centrifuged to pellet the cells. The cell pellets were resuspended in 40% Percoll and layered onto a discontinuous Percoll gradient. Centrifugation at 80% Percoll allowed for the separation of cells at the 40–80% Percoll interface, which were collected as CNS mononuclear cells. The collected CNS mononuclear cells were washed with PBS to remove any remaining Percoll. CNS mononuclear cells were stimulated for 4 hours with PMA and ionomycin in the presence of Brefeldin A to induce cytokine production. After stimulation, cells were fixed, rendered permeable, and stained with appropriate antibodies for intracellular cytokine detection.

### Histological analyses

The lungs, skin, liver, and colon were excised from three-week-old mice. Prior to histological analysis, the samples were fixed in formalin, embedded in paraffin, and stained with hematoxylin and eosin. For CNS histology, spinal cords were fixed in 4% paraformaldehyde, paraffin-embedded, sectioned, and stained with Luxol Fast Blue and hematoxylin and eosin (H&E). To examine colon histology, colons from *Rag1^−/−^* hosts were similarly processed and stained with H&E.

### Adoptive transfer colitis model

Colitis was induced following the protocol described^25^. In brief, CD4^+^ YFP^+^ T_reg_ cells were isolated from 3-week-old CD45.2^+^*Foxp3*^cre^*Cxxc1*^fl/fl^ and CD45.2^+^ *Foxp3*^cre^ mice. A total of 2×10^5^ T_reg_ cells from each group were mixed with 4×10^5^ T_eff_ cells (CD45.1^+^CD4^+^CD45RB^hi^) sorted from CD45.1^+^ mice and transferred into the *Rag1^−/-^* mice via intraperitoneal injection. T_eff_ cells alone were transferred as a control group. Mouse body weight was measured weekly post-adoptive transfer. The percentage change in body weight was calculated by comparing the current weight with the initial weight on day 0. Mice were euthanized when any had reached 80% of their initial body weight. The large intestines were sectioned into 4μm thick slices and stained with hematoxylin.

### In vitro T_reg_ suppression assay

Naïve CD4^+^ T cells isolated from WT mice were labeled with CFSE (C34554; Invitrogen). CD4^+^YFP^+^ T_reg_ cells from *Foxp3*^cre^ and *Foxp3*^cre^*Cxxc1*^fl/fl^ mice were cultured with naïve CD4^+^ T cells (1C10^5^ cells) at various ratios in the presence of 2μg/mL anti-CD3(16-0031-85; Invitrogen) and 3μg/mL anti-CD28(16-0281-85; Invitrogen). On day 3, cells were analyzed by flow cytometry.

### CUT&Tag

CUT&Tag assays of CD4^+^YFP^+^ T_reg_ cells were conducted as previously described^26^. Briefly, approximately 1×10^5^ single cells were carefully pipetted into wash buffer twice. The pelleted cells were resuspended in wash buffer, activated concanavalin (BP531; Bangs Laboratories), and incubated for 15 minutes at room temperature. Cells bound to the beads were resuspended in Dig-Wash Buffer and incubated with a 1:50 dilution of primary antibodies (rabbit anti-H3K4me3, Active Motif,39016; rabbit anti-CXXC1, abcam, ab198977; rabbit anti-FOXP3, abcam, ab 150743;normal IgG, Cell Signaling, 2729) at 4 C overnight. The beads were incubated with a secondary antibody (goat anti-rabbit IgG; SAB3700883; Sigma-Aldrish) diluted 1:100 in Dig-Wash buffer for 60 minutes at room temperature. Cells were treated with Hyperactive pG-Tn5 Transposase (S602; Vazyme) diluted in Dig-300 Buffer for 1 hour at room temperature. The cells were subsequently resuspended in Tagmentation buffer (10mM MgCl2 in Dig-300 Buffer) and incubated at 37C for 1 hour. To halt tagmentation, 10μl was spiked with 0.5M EDTA, 3μl with 10%SDS, and 3μl with 20mg/ml Proteinase K and incubated at 55C for 1h. DNA library amplification was performed according to the manufacturer’s instructions and purified using VAHTS DNA Clean Beads (N411; Vazyme). Libraries were sequenced on the Illumina NovaSeq platform ( Annoroad Gene Technology).

### CUT&Tag and ChIP-seq data analysis

FOXP3 ChIP-seq data was obtained from GSE121279. H3K27me3 ChIP-seq data was obtained from GSE14254. CUT&Tag and ChIP-seq reads were trimmed to 50 bp and aligned against the mouse genome build mm9 using Bowtie2(v2.3.4.1) with default parameters. All PCR duplicates and unmapped reads were removed. Peak calling was performed using MACS2(v2.1.1.20160309) and signal tracks for each sample were generated using the ‘wigToBigWig’ utility of UCSC. We classified the H3k4me3 peaks around TSSs into three groups: broad (>5 kb), medium (1-5kb), and narrow (<1 kb). The top 5% of the widest peaks were considered as broad peaks. The average intensity profiles were generated using deepTools (v2.5.4). Motif analysis was performed using the ‘findmotifsGenome.pl’ command inHomer2 package. Epigenetic factors were identified using the Epigenetic Factor Database(https://epifactors.autosome.org/) and then screened for those that exclusively regulate the expression of their target genes by modulating the deposition of H3K4me3. Genomic distribution was analyzed using the “genomation” R package. GO pathway analysis was performed using the “clusterProfiler” R package. The sequencing information of CUT&Tag data used in this study is summarized in Supplementary Table S1.

### Clustering analysis

Promoters were defined as ±2 kb regions flanking the annotated TSS. Reads in promoters were counted using the “coverage” command in bedtools (v2.26.0) and further normalized to RPKM. The k-means clustering of H3K4me3 and H3K27me3 enrichment at promoters was conducted using the “kmeans” function in R.

### RNA-seq and data analysis

Total RNA was extracted from sorted CD4CYFPC T_reg_ cells using the RNeasy Plus Mini Kit (Qiagen, #74134), following the manufacturer’s protocol. RNA-Seq libraries were constructed and sequenced by Haplox (Nanchang, China), using an Illumina platform with paired-end reads of 150 bp. RNA-seq data was obtained from GSE82076. Raw reads were trimmed to 50 bp and mapped to the mouse genome (mm9) using TopHat (v2.1.1) with default parameters. Only uniquely mapped reads were kept for downstream analysis. The RNA abundance of each gene was quantified using Cufflinks (v2.2.1).

### Single-Cell RNA-Sequencing

A total of 300,000 sort-purified CD4^+^YFP^+^ T_reg_ cells from *Foxp3*^cre^ and *Foxp3*^cre^ *Cxxc1*^fl/fl^ were resuspended in BD Pharmingen Stain Buffer (FBS) (554656; BD). Single cells were isolated using a chromium controller (BD platform, BD Bioscience) according to the manufacturer’s instructions, as previously described^27^. The single cells were labeled with sample tags using the BD Mouse Immune Single-Cell Multiplexing Kit (633793; BD). Following standard protocols, cDNA amplification and library construction were performed to generate scRNA-seq libraries.

### Targeted scRNA-seq data processing

The raw FASTQ files were processed by BD Rhapsody using the Targeted analysis pipeline. After alignment and filtering, the distribution-based error correction(DBEC)-adjusted molecules were loaded into R studio(version 4.3.2). All subsequent analyses were performed using the package Seurat (version 4.4.0) with default parameters. Specifically, the scRNA-seq data counts were log-normalized. All targeted genes were scaled and then were used for Principal components analysis (PCA). The batch effects were removed by the HarmonyMatrix function in the Harmony package (version 1.1.0). The first 20 principal components were used to calculate nonlinear dimensionality reduction using RunUMAP. Differential gene expression (DEGs) between clusters was assessed using the FindAllMarkers function. The clusters were then annotated based on DEGs. Barplots were generated using ggplot2 (version 3.4.4). Heatmaps were generated using pheatmap (version 1.0.12).

### Analysis of the single-cell TCR-seq repertoire

Raw V(D)J fastq reads were processed using BD Rhapsody Pipeline and then were analyzed using scRepertoire (version 1.12.0). The TCR clonotype was called using the nucleotide sequence of the CDR3 region for both TCR alpha and beta chains. For cells with multiple chains, the top two clonotypes with the highest expression were selected for downstream analysis. Clonal overlap between different cell types was calculated using the clonalOverlap function of scRepertoire. A clonotype was defined as expansion if it could be detected in at least two cells.

### scRNA-seq trajectory analysis

UMAP embeddings obtained from the Seurat package were projected into the Slingshot(version 2.10.0) package to construct pseudotime Trajectories for T_reg_ cells. Naïve subsets were set as the root state.

### Whole genome bisulfite sequencing (WGBS) and data analysis

Sorted CD4^+^YFP^+^ T_reg_ cells(3C10^6^) were lysed in cell lysis buffer to release DNA. The bisulfite-treated DNA was used to prepare the sequencing library. DNA libraries were transferred to the Illumina Platform for sequencing using 150 bp paired-end reads. Raw reads were trimmed using TrimGalore (v0.4.4) with default parameters. Subsequently, the reads were mapped against the mm9 reference genome using Bismark v0.19.0 with parameters ‘--bowtie2’. PCR duplicates were removed and the methylation levels were calculated using ‘bismark_methylation_extractor’. We calculated the mean CpG methylation levels of various genome elements: promoter, 5′-UTR, exon, intron, 3′-UTR, genebody, intergenic, CGIs, and repeats using in-house scripts. The sequencing information of WGBS data used in this study is summarized in Supplementary Table S2.

### Statistical analysis

The statistical significance analysis was performed using Prism 8.0 (GraphPad). Error bars are presented as mean ± SD. *P* values of < 0.05 were deemed statistically significant (\**P* < 0.05, \*\**P* < 0.01, \*\*\**P* < 0.001 and \*\*\*\**P* < 0.000). Statistical analyses were performed with unpaired Student’s *t*-test or two-way ANOVA and Holm–Sidak post hoc test.

## Results

### FOXP3 binds regulatory loci primed for activation and repression in T_reg_ cells

FOXP3-mediated gene expression is well-recognized, with several studies highlighting its dual role as both a transcriptional activator and repressor ^28–32^. However, the mechanisms by which FOXP3 regulates T_reg_-specific gene transcription via epigenetic modifications remain incompletely understood. To investigate these mechanisms, we employed CUT&Tag to generate genome-wide H3K4me3 maps in T_reg_ cells. To complement this, we compared our data with an H3K27me3 ChIP-Seq dataset from Wei *et al.*^33^ focusing on previously identified FOXP3-bound loci(Konopacki *et al.* 2019)^34^. This integrative analysis allowed us to identify FOXP3-dependent genes associated with either H3K4me3 (indicative of transcriptional activation) or H3K27me3 (indicative of repression) deposition. These findings provide insight into how FOXP3 modulates the epigenetic landscape to regulate T_reg_ cell function. As expected, H3K4me3 was enriched at gene promoters (fig. S1, A and B). A Venn diagram revealed overlap between FOXP3 binding sites and H3K4me3 peaks, with minimal overlap with H3K27me3 peaks (Fig. 1, A and B). The overlapping regions between FOXP3 binding sites and H3K4me3 or H3K27me3 peaks were predominantly located at promoters (Fig. 1, A and B). To further investigate the role of FOXP3 in promoting H3K4me3 deposition, we compared H3K4me3 levels between FOXP3-positive T_reg_ cells and FOXP3-negative conventional T cells (Tconv). Consistent with our hypothesis, H3K4me3 abundance was higher at T_reg_-specific gene loci (e.g., *Tnfrsf18*, *Ctla4*, *Il2ra* and *Nt5e*) in T_reg_ cells compared to Tconv cells (Fig. 1,C and fig. S1, C). Notably, the selection of these loci was guided by prior studies identifying genes specifically associated with T_reg_ cell function^35^. This finding highlights the role of FOXP3 in shaping the epigenetic landscape of T_reg_ cells through enhanced H3K4me3 deposition at these critical loci.

**Fig. 1.**
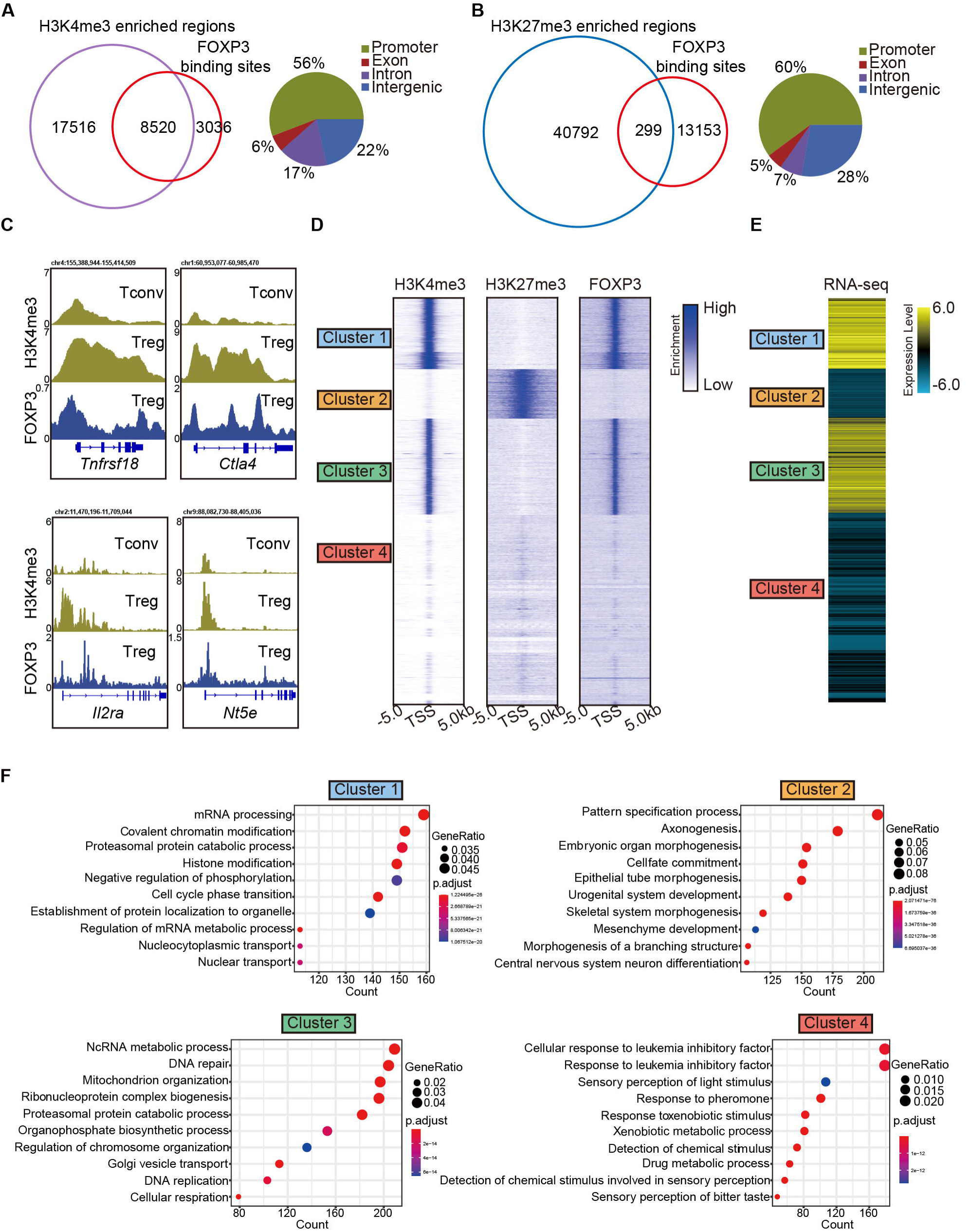
H3K4me3 is required for FOXP3-dependent gene activation in T_reg_ cells. (**A**) Venn diagram showing overlap of H3K4me3 enriched regions (this study) and FOXP3 binding sites (Konopacki *et al*.2019) in sorted CD4^+^YFP^+^T_reg_ cells (left). Genomic distribution of overlapped peaks(right). Note that the overlapped peaks are predominantly enriched at promoters. (**B**) Venn diagram showing overlap of H3K27me3 enriched regions (Wei *et al*.2009) and FOXP3 binding sites in T_reg_ cells (left). Genomic distribution of overlapped peaks (right). Note that the overlapped peaks are predominantly enriched at promoters. (**C**) Representative genome browser view showing the enrichments of H3K4me3 and FOXP3 inTconv or T_reg_ cells. (**D**) Heatmap showing enrichment of H3K27me3, H3K4me3, and FOXP3 surrounding the TSS. Unsupervised K-means clustering was conducted on H3K27me3 and H3K4me3 signals. (**E**)Heatmap showing gene expression levels in T_reg_ cells (RNA-seq data was obtained from Oh *et al.*2017) The clusters were consistent as in Fig. 1C. (**E**) Gene Ontology (GO) pathway analysis of different clusters.

To further characterize these modifications,we clustered promoters into four groups based on the enrichment of H3K4me3 and H3K27me3. Clusters 1 and 3 showed strong enrichment of H3K4me3; cluster 2 was enriched with H3K27me3; and cluster 4 showed weak enrichment of both modifications. FOXP3 preferentially bound to clusters 1 and 3 prom-oters, which displayed high H3K4me3 levels (Fig. 1C). Correspondingly, genes in these clusters exhibited high transcription levels, as shown by reanalysis of previously published RNA-seq data (Oh *et al.* 2017) ^36^ (Figure 1D, right). In contrast, genes with H3K27me3 enrichment at their promoters were transcribed at low levels.

Gene Ontology (GO) analysis of these four clusters revealed distinct functional roles. Cluster 1 was enriched in genes involved in mRNA processing, covalent chromatin modification, and histone modification, while cluster 3 was enriched in genes related to DNA repair and mitochondrion organization (Fig. 1E). Cluster 2, enriched with H3K27me3, was associated with pattern specification process, while cluster 4 showed no correlation with T_reg_ cells. Notably, signature T_reg_ cell genes such as *Tnfrsf18*, *Nrp1*, *Stat5a*, *Lag3*, *Icos*, and *Pdcd1* were enriched in clusters 1 and 3, showing strong H3K4me3 marks (fig. S1C). Conversely, genes like *Hic1*, *Trp73,* and *Rnf157*, associated with inflammatory responses, were enriched for H3K27me3 in clusters 2 (fig. S1C). These findings collectively support the conclusion that FOXP3 contributes to transcriptional activation in T_reg_ cells by promoting H3K4me3 deposition at target loci, while also regulating gene expression directly or indirectly through other epigenetic modifications.

### CXXC1 interacts with FOXP3 and binds H3K4me3-enriched sites in T_reg_ cells

We conducted an enrichment analysis of known motifs at the overlapping peaks of FOXP3 ChIP-seq and H3K4me3 CUT&Tag in T_reg_ cells to identify epigenetic factors that directly interact with FOXP3 to mediate chromatin remodeling and transcriptional reprogramming. Motif analysis of the overlapping peaks between FOXP3 binding sites and regions enriched in H3K4me3 revealed that, in addition to transcription factors, the most abundant motif associated with H3K4me3 was the epigenetic factor CXXC1(fig. S2A). To investigate this further, we performed CUT&Tag for endogenous CXXC1 in T_reg_ cells to examine the genome-wide co-occupancy of CXXC1 and FOXP3. Over half of these CXXC1 binding sites were located at promoter regions (fig. S2B). Additionally, CXXC1 exhibited strong binding at transcription start site (TSS) and CpG islands (CGIs)( Fig. 2A and fig. S2C). As illustrated by the Venn diagram (Fig. 2B), more than half of the FOXP3-bound genes and H3K4me3-enriched genes were also bound by CXXC1. Similarly, more than half of CXXC1 peaks were overlapped with FOXP3 peaks (fig. S2D). Furthermore, the CXXC1-specific and FOXP3-specific binding sites also demonstrated modest binding of FOXP3 and CXXC1, respectively (Fig. 2C). These findings indicate that FOXP3 and CXXC1 share a substantial number of target genes in T_reg_ cells. To confirm this interaction, we further validated the reciprocal immunoprecipitation of both endogenous and exogenous CXXC1 and FOXP3 (Fig. 2D and fig. S2E). An immunofluorescence assay revealed predominant colocalization of CXXC1 with FOXP3 in the nucleus (Fig. 2E). Overall, these results suggest that CXXC1 primarily functions as a coactivator of FOXP3-driven transcription in T_reg_ cells.

**Fig. 2.**
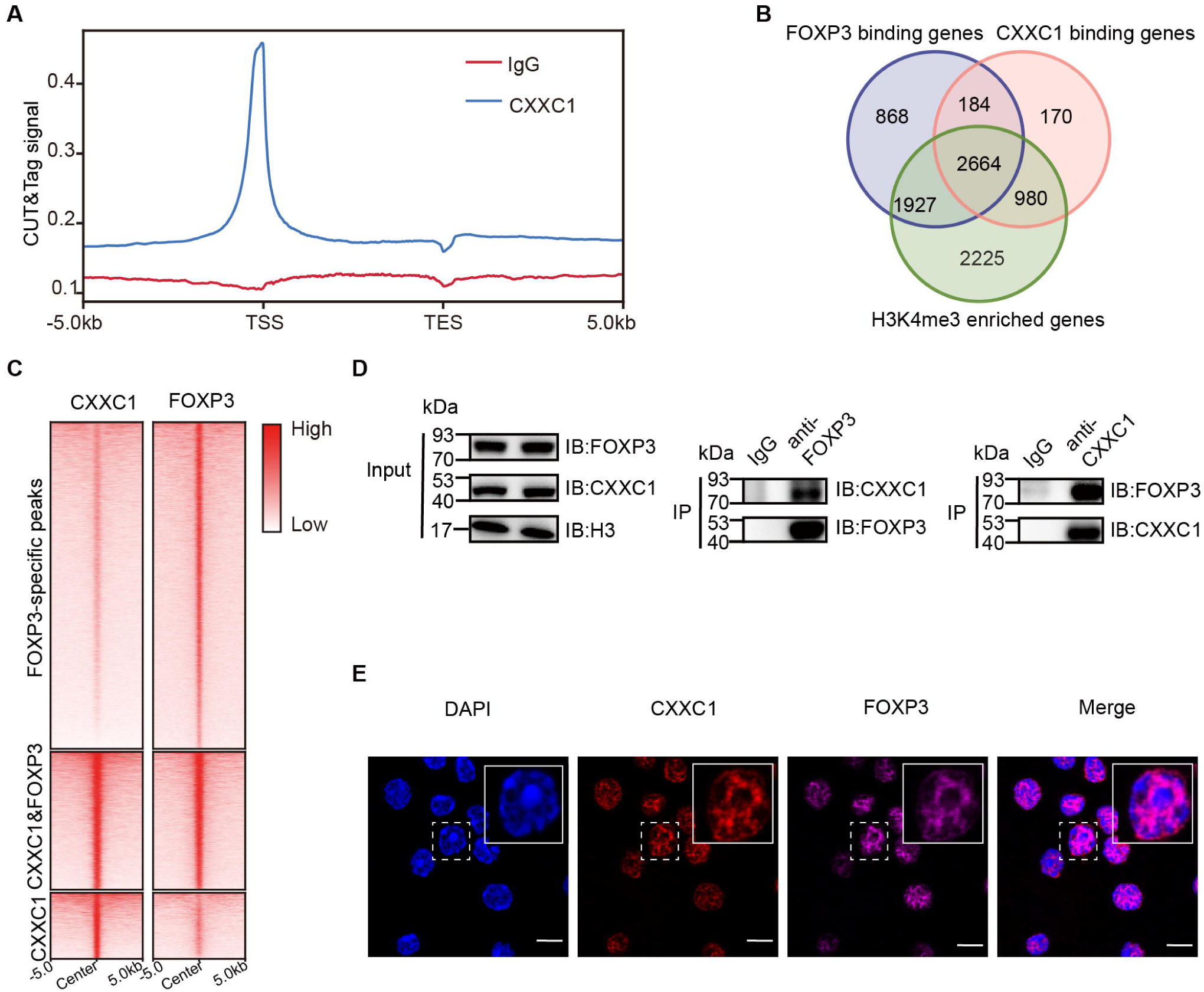
CXXC1 interacts with FOXP3 in T_reg_ cell. (**A**) Average CXXC1 CUT&Tag signals around promoters in T_reg_cells. IgG was used as the control. (**B**) Venn diagrams showing the overlap of FOXP3 binding genes, CXXC1 binding genes, and H3K4me3 enriched genes in T_reg_ cells. Gene promoters covered by FOXP3 binding sites, CXXC1 binding sites, or exhibited high H3K4me3 levels were defined as FOXP3-bound genes, CXXC1-bound genes, or H3K4me3-enriched genes. (**C**) Heatmaps showing FOXP3 ChIP-seq and CXXC1 CUT&Tag signals at indicated regions. (**D**) Interaction between FOXP3 and CXXC1 was assessed by co-IP (forward and reverse) using T_reg_ cell lysates. (**E**) Immunofluorescence for FOXP3 and CXXC1 colocalization in T_reg_ cells. Scale bars, 2 μm.

### Complete ablation of *Cxxc1* in T_reg_ cells leads to a fatal autoimmune disease

To investigate the role of CXXC1 in T_reg_-cell homeostasis and function, we generated *Foxp3*^Cre^*Cxxc1*^fl/fl^ mice (conditional knockout [cKO] mice) by crossing *Cxxc1*^fl/fl^ with *Foxp3*^YFP-Cre 37^ mice, thereby specifically deleting *Cxxc1* in T_reg_ cells. The effective depletion of *Cxxc1* in T_reg_ cells was confirmed through quantitative PCR (qPCR) and Western blotting (fig. S3A). Notably, cKO mice appeared normal at birth but later exhibited spontaneous mortality starting around three weeks of age (Fig. 3A). Deletion of *Cxxc1* in T_reg_ cells led to the development of severe inflammatory disease, characterized by reduced body size, stooped posture, crusting of the eyelids, ears, and tail, and skin ulceration, particularly on the head and upper back (Fig. 3, B and C). Additionally, cKO mice developed extensive splenomegaly and lymphadenopathy (Fig. 3D). *Foxp3*^Cre^*Cxxc1*^fl/fl^ mice exhibited elevated serum levels of anti-dsDNA autoantibodies and IgG , along with a modest increase in IgE concentration (Figure 3E and fig. S3B). Histopathological analysis revealed massive lymphocyte and myeloid cell infiltration in the skin, lungs, liver sinusoids, and colon mucosa (Fig. 3F). In full agreement with the aforementioned severe autoimmune diseases, *Foxp3*^Cre^*Cxxc1*^fl/fl^ mice had decreased percentages and numbers of CD4^+^ Foxp3^+^ T_reg_ cells in small intestine lamina propria (LPL), liver and lung(Fig. 3G and fig. S3C). Moreover, cKO mice displayed an increase in CD8^+^ T cell percentages (fig. S3D), along with a marked rise in cells exhibiting an effector/memory phenotype (CD44hi CD62Llo) (Fig. 3H). Furthermore, T cells from cKO mice produced elevated levels of IFN-γ, IL-17, and IL-4 in CD4^+^ YFP^−^ T cells, as well as increased IFN-γ production in CD8^+^ T cells (Fig. 3I and fig. S3, E and F). These phenotypes closely resembled those observed in *Foxp3-*deficient mice ^38^ or mice with depleted T_reg_ cells^39^, suggesting a deficiency in immune suppression.

**Fig. 3.**
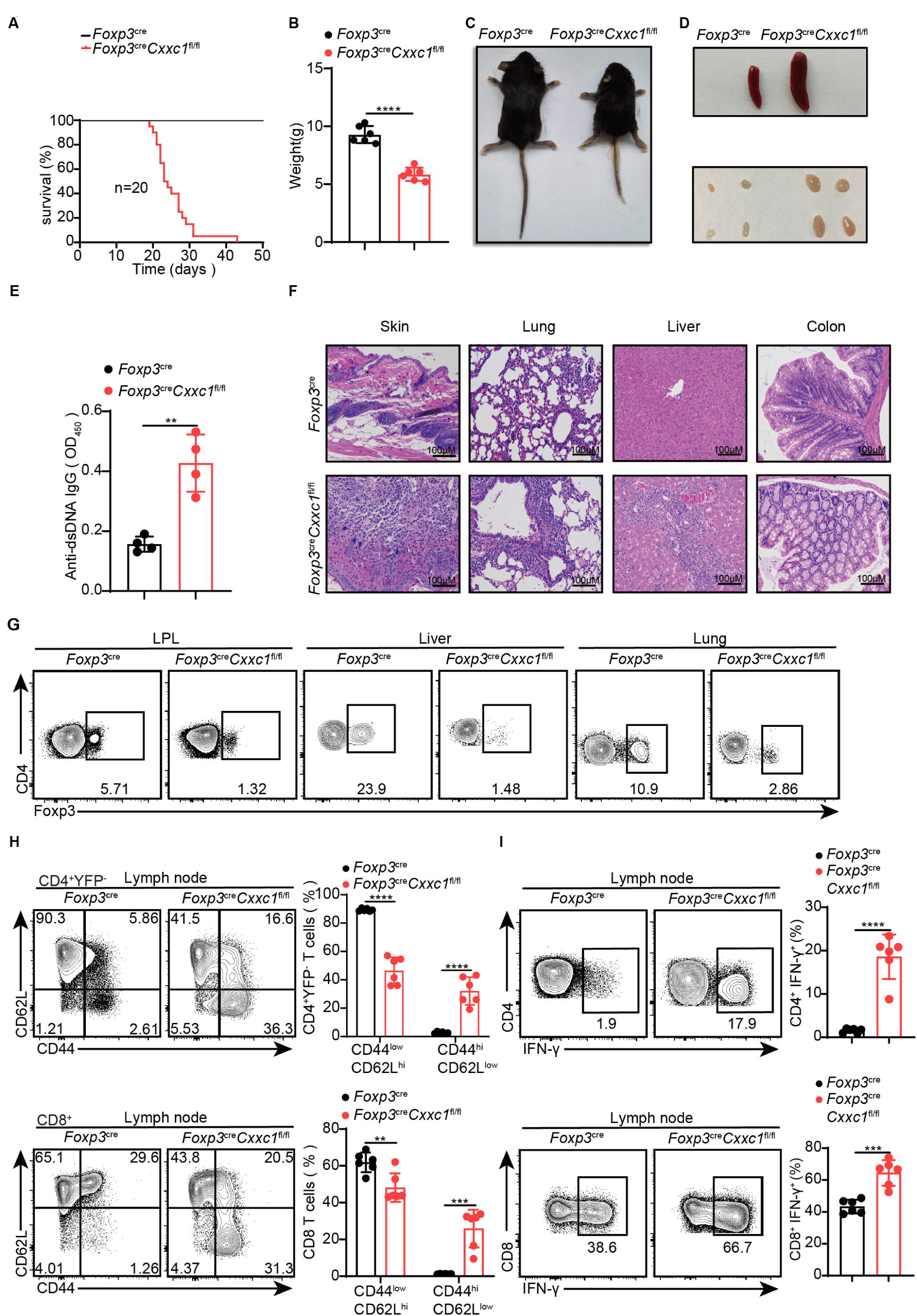
*Foxp3creCxxc1*^fl/fl^ mice spontaneously develop a fatal early-onset inflammatory disorder. (**A**) Survival curves of *Foxp3*^cre^ (black line) and *Foxp3*^cre^*Cxxc1*^fl/fl^ (red line) mice (*n* = 20). (**B**) Gross body weight of *Foxp3*^cre^ and *Foxp3*^cre^*Cxxc1*^fl/fl^ mice (*n*C=C6). (**C**) A representative image of *Foxp3*^cre^ and *Foxp3*^cre^*Cxxc1*^fl/fl^ mice. (**D**) Representative images showing the spleen and peripheral lymph nodes from *Foxp3*^cre^ and *Foxp3*^cre^*Cxxc1*^fl/fl^ mice. (**E**) ELISA quantification of anti-dsDNA IgG in the serum of *Foxp3*^cre^ and *Foxp3*^cre^*Cxxc1*^fl/fl^ mice (n = 4). (**F**) Hematoxylin and eosin staining of the skin, lung, liver, and colon from *Foxp3*^cre^ an d *Foxp3*^cre^*Cxxc1*^fl/fl^ mice (scale bar, 100Cμm). (**G**) Representative flow cytometry plots of CD4C Foxp3C T_reg_ cells isolated from the small intestinal lamina propria (LPL), liver, and lung of *Foxp3*^cre^ and *Foxp3*^cre^*Cxxc1*^fl/fl^ mice. (**H**) Flow cytometry analysis of CD62L and CD44 expression on peripheral lymph node CD4^+^YFP^−^ and CD8^+^T cells from *Foxp3*^cre^ and *Foxp3*^cre^*Cxxc1*^fl/fl^ mice (left). Right, frequency of CD44^low^CD62L^hi^ and CD44^hi^CD62L^low^ population in CD4^+^ YFP^−^ or CD8^+^ T cells (*n* = 6). (**I**) Lymph node cells from *Foxp3*^cre^ and *Foxp3*^cre^*Cxxc1*^fl/fl^ mice were stimulated ex vivo with PMA + ionomycin for 4h and analyzed for IFN-γ expressing in CD4^+^ YFP^−^ or CD8^+^ T cells using flow cytometry (left). Right, percentages of IFN-γ^+^CD4^+^ YFP^−^ or IFN-γ^+^CD8^+^ T cells in the lymph nodes of *Foxp3*^cre^ and *Foxp3*^cre^*Cxxc1*^fl/fl^ mice (*n* = 6). All mice analyzed were 18-20 days old unless otherwise specified. Error bars show mean ± SD. The log-rank survival curve was used for survival analysis in A, and multiple unpaired *t*-test or two-tailed Student’s *t*-test were used for statistical analyses in B, E, Gand H (\*\**P*<0.01, \*\*\**P*C<C0.001, \*\*\*\**P*<0.0001). The flow cytometry results are representative of three independent experiments.

### CXXC1 is necessary for the maintenance of T_reg_-cell suppressive activity

Despite the development of severe autoimmune disease, we observed an increase in both the absolute number and percentage of FOXP3^+^ T_reg_ cells in the lymph nodes (fig. S4A). The expression level of the FOXP3 protein was only slightly altered in *Cxxc1*-deficient T_reg_ cells (fig. S4B). In an *in vitro* suppression assay, T_reg_ cells from *Foxp3*^Cre^*Cxxc1*^fl/fl^ and WT mice exhibited similar suppressive effects on naïve T (Tn) cell proliferation (fig. S4C). The expression of the hallmark T_reg_-cell marker CTLA-4 showed a modest increase in *Cxxc1*-deficient T_reg_ cells compared to WT T_reg_ cells, while the expression of GITR remained unchanged (fig. S4D). To further assess the suppressive capacity of *Cxxc1*-deficient T_reg_ cells *in vivo*, we employed the experimental autoimmune encephalomyelitis (EAE) model. Naïve CD4^+^ T cells from 2D2 mice were co-transferred with T_reg_ cells from either *Foxp3*^Cre^ or *Foxp3*^Cre^ *Cxxc1*^fl/fl^ mice into *Rag1^−/-^* recipients, and EAE was induced in these recipient mice. Mice that received only naïve CD4^+^ T cells from 2D2 mice developed more severe EAE symptoms (Fig. 4A). The addition of WT T_reg_ cells from *Foxp3*^Cre^ mice slightly mitigated EAE progression and reduced Th17 cells in the spinal cord (Fig. 4, A to D). In contrast, *Foxp3*^Cre^ *Cxxc1*^fl/fl^ T_reg_ cells failed to suppress EAE (Fig. 4, A to D), and the cKO mice showed a reduction in T_reg_ cells frequency in CNS tissues(Fig. 4E). Finally, we examined the role of CXXC1 in T_reg_ cell-mediated suppression using T cell transfer-induced colitis, in which naïve T cells were transferred to *Rag1^−/-^* recipients either alone or together with WT or *Foxp3*^Cre^ *Cxxc1*^fl/fl^ T_reg_ cells. The transfer of naïve T cells led to weight loss and intestinal pathology in recipient mice(Fig. 4, F and G). Mice receiving WT T_reg_ cells continued to gain weight (Fig. 4F) , whereas those that received T_reg_ cells from cKO mice were unable to prevent colitis and exhibited a reduced percentage of T_reg_ cells (Fig. 4, F to H). These findings underscore the critical role of CXXC1 in maintaining T_reg_ cell function *in vivo*.

**Fig. 4.**
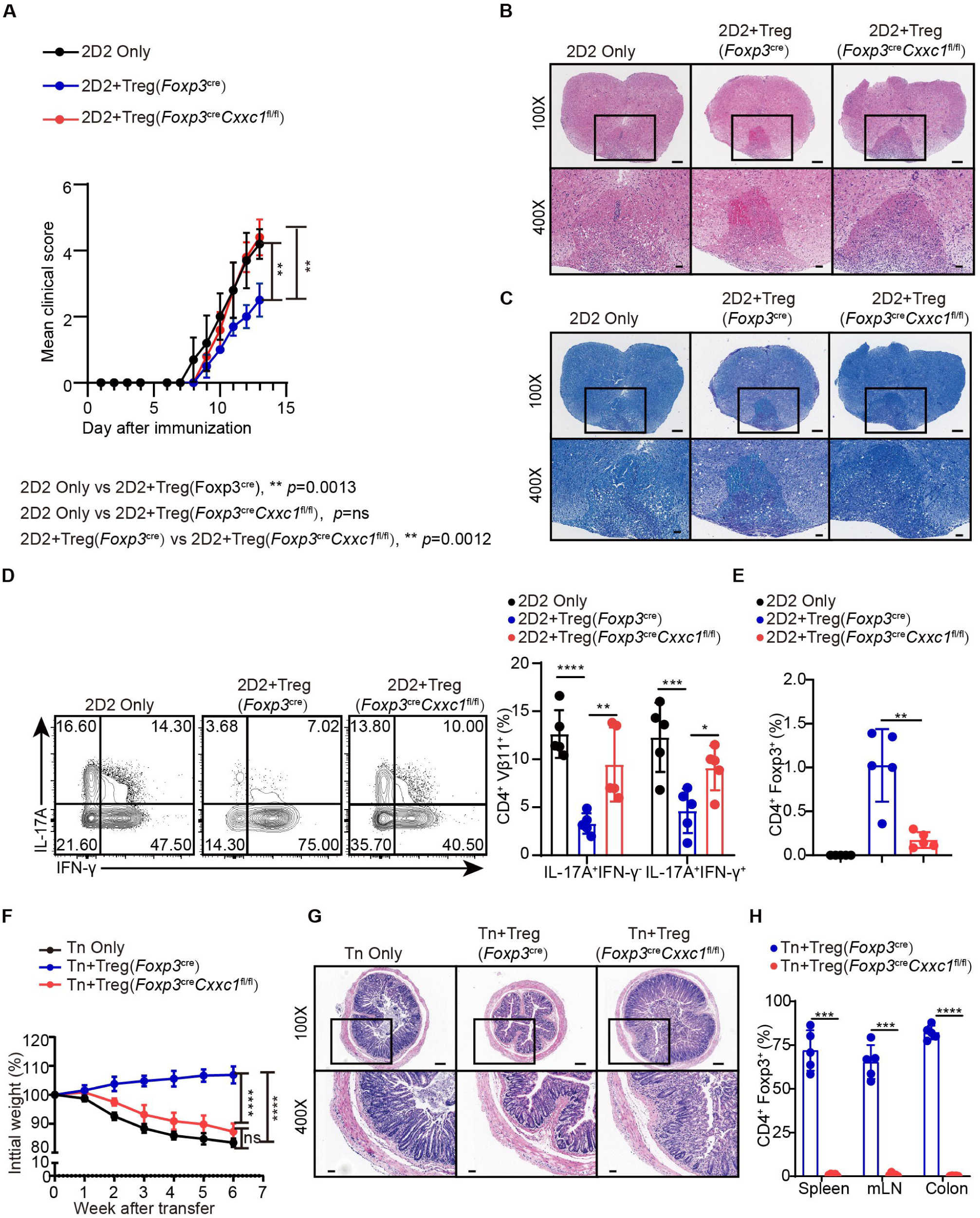
CXXC1 is essential for T_reg_ cells to suppress T cell-mediated EAE and colitis. (**A**)Mean clinical scores for EAE in *Rag1^−/−^* recipients of 2D2 CD4^+^ T cells, either alone or in combination with *Foxp3*^cre^ or *Foxp3*^cre^*Cxxc1*^fl/fl^ mice after immunization with MOG35–55, complete Freund’s adjuvant (CFA), and pertussis toxin (n=5). (**B** and **C**)Representative histology of the spinal cord of *Rag1^−/−^* mice after EAE induction. Hematoxylin and eosin(H&E) staining (upper), Luxol fast blue (F&B) staining (lower). Scale bars, 50 μm (400×) and 200 μm (100×). (**D**) Representative flow cytometry plots and quantification of the the percentages of IFNγ^+^ or IL-17A^+^ CD4^+^Vβ11^+^ T cells (n=5). (**E**) Statistical analysis of the percentage CD4^+^ FOXP3^+^ T_reg_ cell in CNS tissues 14 days after EAE induction. (**F**) Changes in body weight of *Rag1^−/−^* mice after colitis induction(n=6). (**G**) Haematoxylin and eosin (H&E) staining of colons from T cell-induced colitis mice 6 weeks after T cell transfer. Scale bars, 50 μm (400×) and 200 μm (100×). (**H**) Statistical analysis of the percentage CD4^+^ FOXP3^+^ T_reg_ cell in the spleen, mesenteric lymph nodes, and colon 6 weeks after colitis induction. Error bars show mean ± SD. *P* values are determined by a two-tailed Student’s *t*-test or two-way ANOVA and Holm–Sidak post hoc test (A, D, E and H). (\**P*<0.05, \*\**P*<0.01, \*\*\**P*C<C0.001, \*\*\*\**P*<0.0001).

### T_reg_ cell lineage homeostasis and proliferation depend upon CXXC1

T_reg_ cells harbor a diverse T-cell receptor (TCR) repertoire, which likely play a critical role in their immune suppression function ^40–42^. To explore the role of CXXC1 in T_reg_-mediated suppression, we performed single-cell RNA sequencing (scRNA-seq) combined with TCR sequencing (TCR-seq) on CD4^+^YFP^+^ T_reg_ cells isolated from mouse lymph nodes. After quality control and removal of doublets, 18,577 cells were retained for further analysis. Through unsupervised clustering and uniform manifold approximation and projection (UMAP) analysis, we identified eight distinct T_reg_ cell clusters based on the expression of well-characterized markers, with a particular focus on two clusters of activated T_reg_ cells that exhibited higher expression of markers and gene sets relative to naïve T_reg_ cells (Fig. 5A and fig. S5, A to C). A comparison between *Cxxc1*-deficient and WT T_reg_ cells within each cluster revealed a reduction in *Cxxc1*-deficient cells in the naïve subsets, while an increase was observed in the Gzmb^+^ and H2-Eb1^+^ subsets (Fig. 5B). To further elucidate the transition of T_reg_ cells along a dynamic biological timeline, we constructed pseudo-time trajectories using Slingshot ^43^. The pseudo-time gradient depicted a progression from quiescent to activated T_reg_ cells, ultimately encompassing the Gzmb^+^ and H2-Eb1^+^ subsets (Fig. 5C). Given the antigen-specific suppression capabilities of T_reg_ cells ^44,45^, we examined their clonal expansion. The analysis revealed that expanded WT TCR clonotypes (nC≥C2) were predominantly distributed among the Nt5e^+^ subsets, while *Cxxc1*-deficient T_reg_ cells showed expanded clonotypes primarily within the Gzmb^+^ and H2-Eb1^+^ subsets(Fig. 5D and Fig. S5D). TCR sharing analysis indicated clonotype sharing among various clusters of WT T_reg_ cells, suggesting a degree of homogeneity. However, the reduced TCR sharing in *Cxxc1*-deficient T_reg_ cells implies that decreased TCR diversity may impair the suppressive activity of T_reg_ cells (Fig. 5E) ^42^. Furthermore, the *Cxxc1*-deficient group exhibited lower expression of several T_reg_-specific genes associated with suppressive functions, such as*Nt5e*, *Il10*, *Pdcd1*, *Klrg1*, as well as genes that inhibit effector T cells differentiation, including *Sell* and *Tcf7* (Fig. 5F and fig. S5E). Conversely, *Cxxc1*-deficient T_reg_ cells demonstrated elevated expression of *Gzmb*, *Il2ra*, and *Cd69* compared to WT, reflecting a profile indicative of increased activation (Fig. 5, F and G). Additionally, we observed increased expression of genes linked to Th1-type inflammation, such as *Ifng*, *Tbx21,* and *Hif1a*, in *Cxxc1*-deficient T_reg_ cells, likely due to extreme inflammatory conditions (Fig. 5, F to H). The proportion of FOXP3^+^Ki67^+^ T_reg_ cells was lower in cKO mice compared to WT mice (Fig. 5I). These findings underscore the crucial role of CXXC1 in in maintaining T_reg_ cell function and homeostasis.

**Fig. 5.**
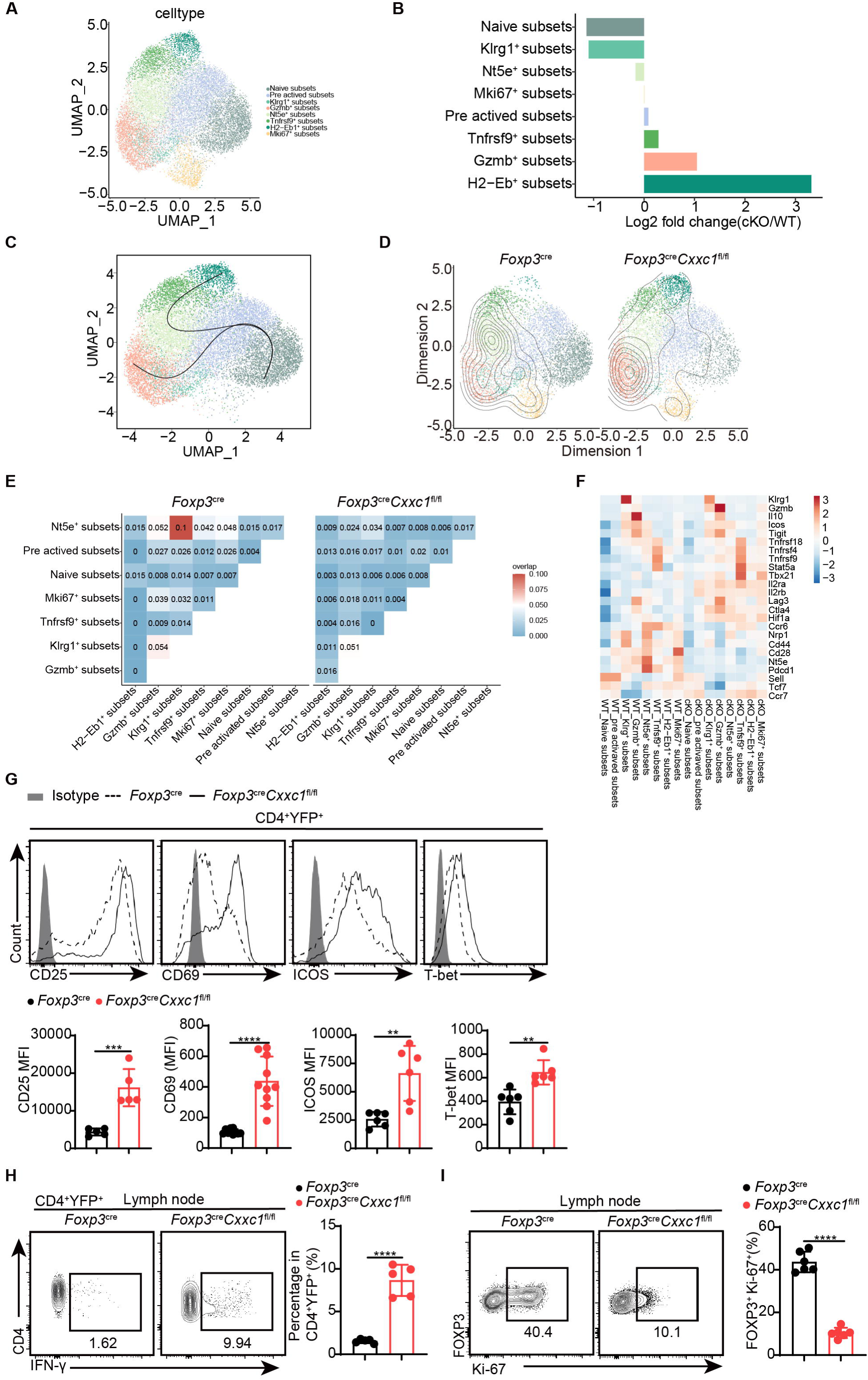
Single-cell transcriptomics reveals distinct T_reg_ cell populations. (**A**) UMAP plot showing clusters identified based on variable gene expression of sorted CD4^+^YFP^+^ T_reg_ cells. Each dot represents a cell, and each color corresponds to a different population of cell types. Clustering analysis revealed 8 distinct T_reg_ cell populations. (**B**) Mean fold changes in cluster abundance between *Foxp3*^cre^ and *Foxp3*^cre^*Cxxc1*^fl/fl^ mice. (**C**)Pseudotime trajectories of T_reg_ cells based on Slingshot, color-coded by T_reg_ cell subpopulations. (**D**) Visualization of density and clonotype richness across T_reg_ clusters from *Foxp3*^cre^ and *Foxp3*^cre^*Cxxc1*^fl/fl^ mice. (**E**) TCR sharing of expanded clonotypes across all possible combinations of T_reg_ cells from *Foxp3*^cre^ and *Foxp3*^cre^*Cxxc1*^fl/fl^ mice. (**F**) Heatmap showing Z scores for the average expression of T_reg_-specific genes in each cluster between *Foxp3*^cre^ and *Foxp3*^cre^*Cxxc1*^fl/fl^ mice. (**G**-**H**)Representative flow cytometry plots and quantification of (G) expression of CD25, CD69, ICOS, T-bet, and (H) IFN-γ in CD4^+^YFP^+^ T_reg_ cells from *Foxp3*^cre^ and *Foxp3*^cre^*Cxxc1*^fl/fl^ mice (n=5 CD25, n=10 CD69, n=5 ICOS, n=6 T-bet, n=5 IFN-γ). (**I**)Ki-67 expression (left) and frequency (right) in CD4^+^FOXP3^+^ T_reg_ cells from *Foxp3* ^cre^ a-nd *Foxp3*^cre^*Cxxc1*^fl/fl^ mice (n=6). Error bars show mean ± SD. *P* values are determined by a two-tailed Student’s *t*-test (G-I). (\*\**P*<0.01, \*\*\**P*□<□0.001, \*\*\*\**P*<0.0001). The flow cytometry results are representative of three independent experiments.

### Intrinsic *Cxxc1* deficiency impairs T_reg_ cell suppression, proliferation, and molecular programs

To confirm that the deficiency in T_reg_ cell function in T_reg_-specific *Cxxc1*-deficient animals is due to intrinsic defects caused by *Cxxc1* deficiency, rather than severe autoimmune inflammation in *Foxp3*^Cre^ *Cxxc1*^fl/fl^ mice, we examined *Cxxc1*-sufficient and *Cxxc1*-deficient T_reg_ cell subsets in heterozygous *Foxp3*^Cre/+^ *Cxxc1*^fl/fl^ (designated as “het-KO”) and littermate *Foxp3*^Cre/+^ *Cxxc1*^fl/+^ (designated as “het-WT”) female mice (Fig. 6A). Notably, het-KO female mice did not exhibit overt signs of autoimmunity, as random X-chromosome inactivation led to the coexistence of both *Cxxc1*-cKO and *Cxxc1*-WT T_reg_ cells. However, both the frequency and absolute numbers of FOXP3^+^YFP^+^ T_reg_ cells within the total T_reg_ population were reduced in het-KO mice compared to their counterparts in het-WT littermates, indicating that *Cxxc1* deficiency imposes a competitive disadvantage on T_reg_ cells (Fig. 6B). Additionally, *Cxxc1*-deficient YFP^+^ T_reg_ cells failed to upregulate the proliferation marker Ki-67 (Fig. 6C). Moreover, YFP^+^ T_reg_ cells in het-KO female mice showed reduced expression of key genes essential for suppressive function, including ICOS, CD25,CTLA4, and GITR, compared to YFP^−^T_reg_ cells from the same mice (Fig. 6D). Consistently, we confirmed the impaired suppressive function of T_reg_ cells from heterozygous *Foxp3*^Cre/+^ *Cxxc1*^fl/fl^ mice in *vitro* and in *vivo* (Fig. 6E and fig. S6A-E). To investigate the molecular program affected by the deletion of *Cxxc1* in T_reg_ cells, we performed RNA-sequencing (RNA-seq) analysis on CD4^+^YFP^+^ T_reg_ cells isolated from het-WT and het-KO mice. We then conducted a differential gene expression (DGE) analysis based on the RNA-seq data. Among all expressed genes, 865 were upregulated (“Up” genes) and 761 were downregulated (“Down” genes) by ≥1.5-fold in CD4^+^YFP^+^ T_reg_ cells from het-KO mice compared to het-WT mice, with a false discovery rate (FDR)–adjusted *P*-value cutoff of 0.05 (fig. S6F). Gene Ontology (GO) enrichment analysis revealed that the downregulated genes in *Cxxc1*-deficient T_reg_ cells were predominantly enriched in pathways related to the negative regulation of immune system process and regulation of cell−cell adhesion (fig. S6G). The *Cxxc1*-deficient T_reg_ cells also showed reduced expression of several genes associated with T_reg_ cell suppressive function, including *Il10*, *Tigit*, *Lag3*, *Icos*, *Nt5e*(encoding CD73) and *Itgae* (encoding CD103) (fig. S6H). Thus, while YFPC WT T_reg_ cells effectively prevent autoimmunity in het-KO mice, the absence of *Cxxc1* in YFPC T_reg_ cells disrupts key T_reg_ cell marker expression and impairs their suppressive function under steady-state conditions.

**Fig. 6.**
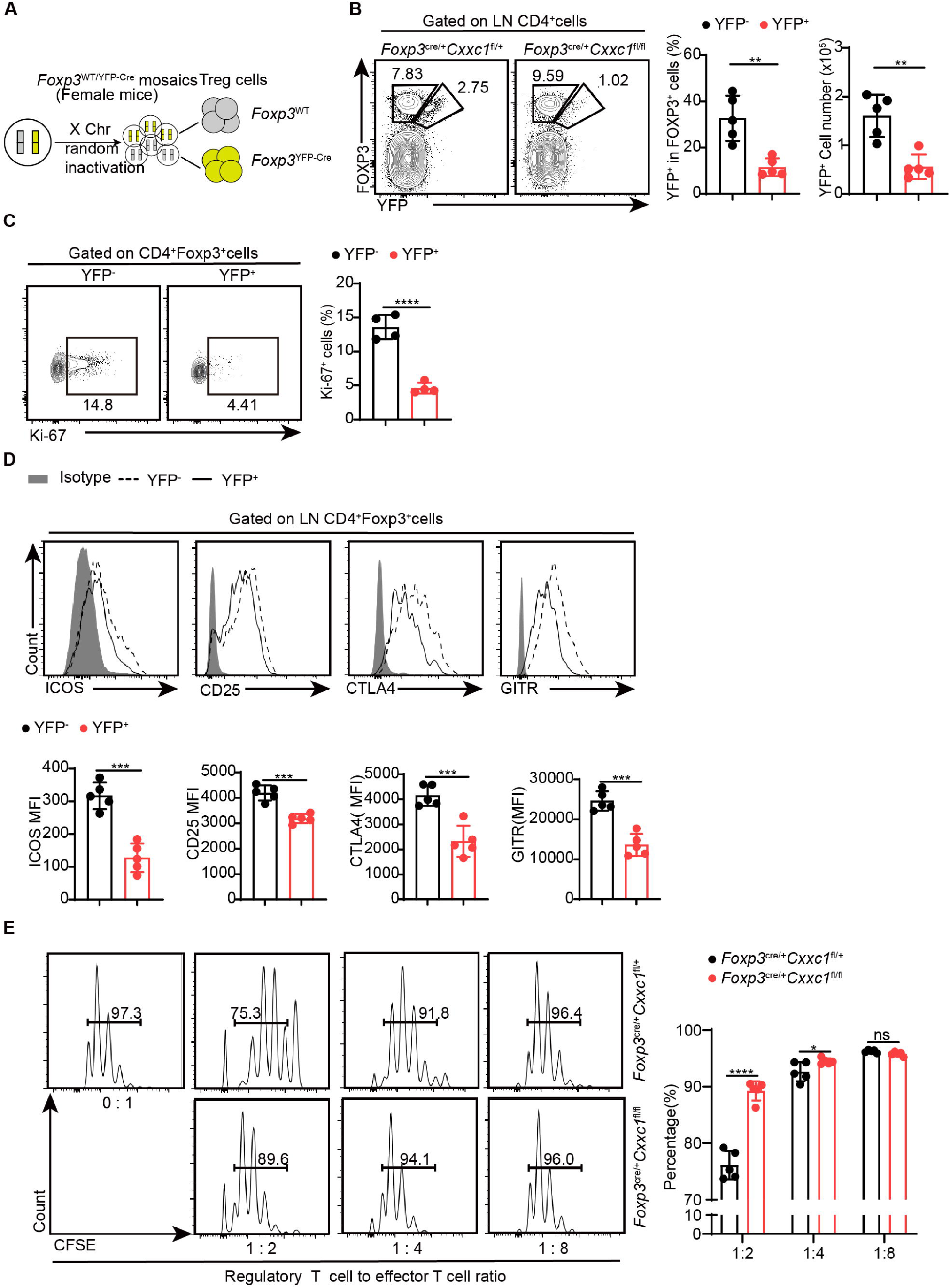
*Cxxc1*-deficient T_reg_ cells exhibit functional impairment and disrupted homeostasis in steady-state conditions. (**A**) Schematic representation of wild-type and Cre-positive T_reg_ cells in female *Foxp3*^Cre/+^mice. (**B**) Flow cytometry analysis of the YFP^−^FOXP3^+^ (WT) and YFP^+^FOXP3^+^ (KO) T_reg_ cells in *Foxp3*^Cre/+^ *Cxxc1*^fl/+^ (het-WT) and *Foxp3*^Cre/+^ *Cxxc1*^fl/fl^ ( het-KO) female mice (left), along with the frequency and absolute numbers of YFP^+^ cells within the total T_reg_ population (right)(n=5). (**C**) Flow cytometry analysis of Ki-67expression (left) and MFI (right) in YFP^−^ and YFP^+^ cells within the CD4^+^FOXP3^+^ T_reg_ cells from 6- to 8-week-old het-KO female mice (n=4). (**D**)Representative flow cytometry plots and quantification of ICOS, CD25, CTLA4, and GITR expression in YFP^−^ and YFP^+^ cells within CD4^+^FOXP3^+^ T_reg_ cells from het-KO female mice (n=5). (**E**)Suppression of CFSE-labelled Tn cell proliferation by different ratios of CD4^+^YFP^+^ T_reg_ cells from *Foxp3*^Cre/+^ *Cxxc1*^fl/+^ and *Foxp3*^Cre/+^ *Cxxc1*^fl/fl^ female mice. On the right, the percentage of proliferated responding T cells is presented (n=5). Error bars show mean ± SD. *P* values are determined by a two-tailed Student’s *t*-test (B-E). (ns, not significant. \**P*<0.05,\*\**P*<0.01, \*\*\**P*C<C0.001, \*\*\*\**P*<0.0001). The flow cytometry results are representative of three independent experiments.

### The FOXP3-CXXC1 complex regulates the expression of key factors in T_reg_ cells that are associated with the breadth of H3K4me3

CXXC1 binds to unmethylated CpG DNA via its N-terminal CXXC finger domain, facilitating its interaction with DNA methyltransferase 1 (DNMT1). This binding stabilizes the DNMT1 protein, thereby regulating DNA methylation ^46,47^. To investigate whether CXXC1 depletion affects DNA methylation in T_reg_ cells, we performed whole genome bisulfite sequencing (WGBS) on T_reg_ cells isolated from both WT and cKO mice. On average, *Cxxc1*-deficient T_reg_ cells exhibited no changes in DNA methylation at gene loci or across genome-wide CpG sites, irrespective of chromosomal region (fig. S7, A to C). Furthermore, *Cxxc1* knockout T_reg_ cells did not show a increase in DNA methylation at key T_reg_ signature gene loci (fig. S7D). Given the pivotal role of MLL4-mediated H3K4me1 in establishing the enhancer landscape and facilitating long-range chromatin interactions during T_reg_ cell development^15^, we performed CUT&Tag to assess changes in H3K4me1 levels in *Cxxc1*-deficient T_reg_ cells. This analysis revealed that H3K4me1 levels were similar in both WT and *Cxxc1*-deficient T_reg_ cells (fig. S7, E to G).

While H3K4me3 modifications typically form sharp 1-to 2-kb peaks around promoters, some genes exhibit broader H3K4me3 regions, referred to as broad H3K4me3 domains (H3K4me3-BDs), which can extend to cover part or all of the gene’s coding sequences (up to 20 kb) ^48,49^. Broad H3K4me3 domains are preferentially associated with genes essential for the identity or function of specific cell types ^48,50^ and have been implicated in enhancing transcriptional elongation and increasing enhancer activity ^50^. To further explore the connection between broad H3K4me3 domains and the expression of immune-regulatory genes, we analyzed genes containing broad H3K4me3 regions. We classified the H3K4me3 domains surrounding transcription start sites (TSSs) into three categories: broad (more than 5 kb), medium (between 1 and 5 kb), and narrow (less than 1 kb) (Fig. 7A). Notably, *Cxxc1*-deficient T_reg_ cells exhibited weaker H3K4me3 signals compared to WT cells within the broad H3K4me3 domains where CXXC1 binding is prominent (Fig. 7, A and B). Using the criteria established by Benayoun *et al.*^48^, which defines the top 5% of the widest H3K4me3 domains as BDs, we observed similar enrichment results (fig. S7, H and I). We then compared three groups of genes: BD-associated genes with reduced H3K4me3 levels following *Cxxc1* deletion, genes with direct CXXC1 binding, and genes with direct FOXP3 binding. The Venn diagram revealed that the majority of genes (283 out of 294, 96%) with CXXC1 binding and reduced H3K4me3 levels overlap with FOXP3-bound genes, suggesting that CXXC1 presence enhances FOXP3 binding within the broad H3K4me3 domain (Fig. 7C). Furthermore, GO term analysis indicated that BD-associated genes are enriched in biological processes related to the negative regulation of immune system processes (Fig. 7D). Genome browser views displayed the enrichments of FOXP3, CXXC1, and H3K4me3 at key signature genes in T_reg_ cells, such as *Ctla4, Il2ra, Icos*, and *Tnfrsf18*, with lower H3K4me3 densities observed at these loci in *Cxxc1*-deficient T_reg_ cells (Fig. 7E). Similar patterns were observed at core genes involved in T_reg_ homeostasis and suppressive function (e.g., *Lag3*, *Nt5e*, *Ikzf4*, and *Cd28*) (fig. S7, J)^51,52^. These findings suggest that CXXC1 and FOXP3 collaboratively promote sustained T_reg_ cell homeostasis and function by preserving the H3K4me3 modification at key T_reg_ cell genes.

**Fig. 7.**
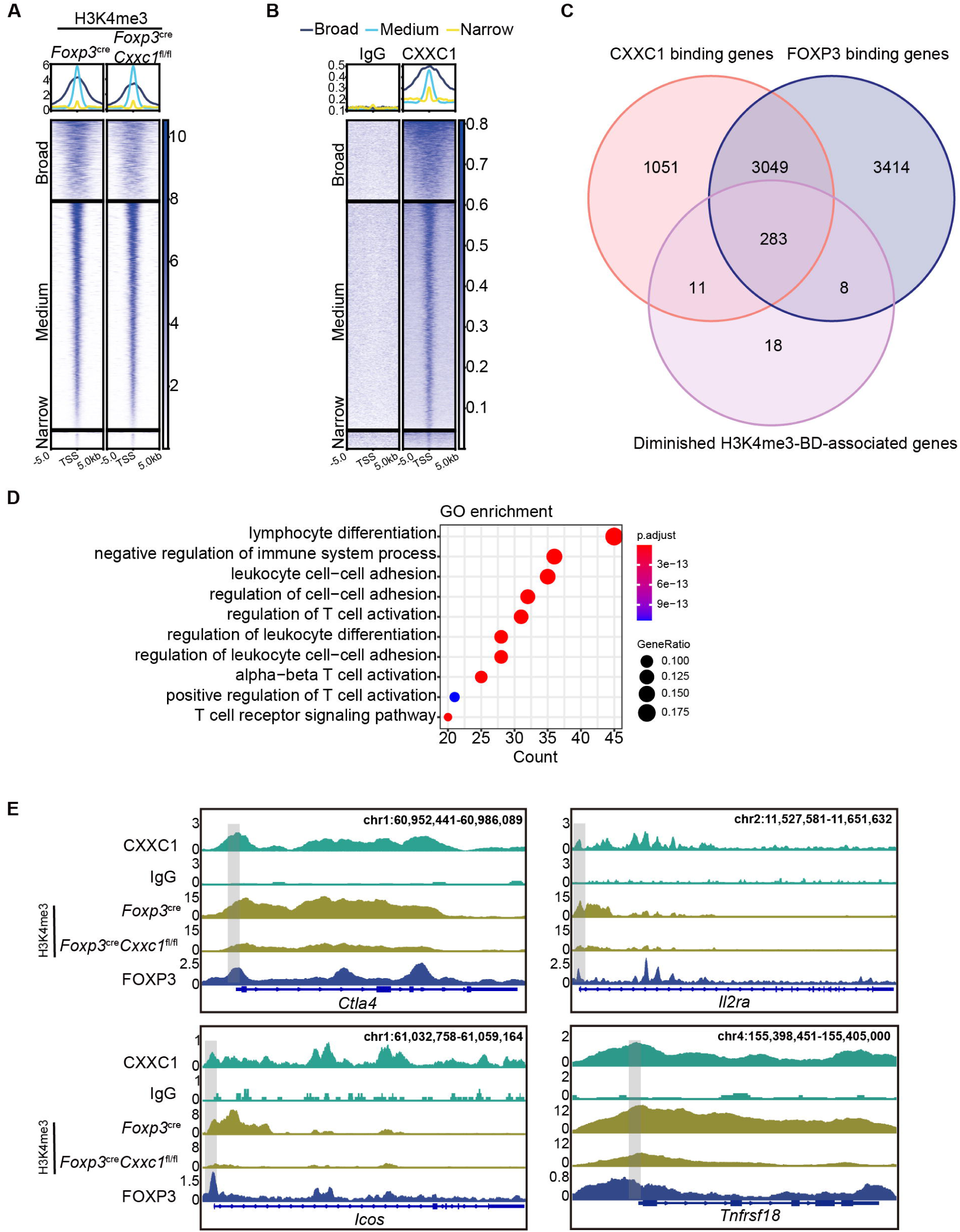
CXXC1 regulates Foxp3-dependent molecule H3K4 trimethylation in T_reg_ cells. (**A**-**B**)Heatmaps showing H3K4me3(A)and CXXC1(B) signals centered on narrow, medium, and broad domains. The top panel shows the average CUT&Tag signals around indicated domains. (**D**) Venn diagrams showing the overlap of FOXP3 binding genes, CXXC1 binding genes, and H3K4me3-BD-associated genes with decreased H3K4me3 levels after CXXC1 depletion in T_reg_ cells. (**E**) Gene Ontology (GO) pathway analysis of the overlap peaks. (**F**) Representative genome browser view showing the enrichments of FOXP3, CXXC1, and H3K4me3 in T_reg_ cells.

We previously demonstrated that the FOXP3-CXXC1 complex plays a key role in modulating H3K4me3 deposition at T_reg_-specific gene loci. To further clarify whether *Cxxc1* deletion affects FOXP3 binding to its target genes, we performed CUT&Tag experiments to compare FOXP3 binding profiles between WT and *Cxxc1* KO T_reg_ cells. The results revealed that most FOXP3-bound regions in WT T_reg_ cells were similarly enriched in KO T_reg_ cells, indicating that *Cxxc1* deletion does not impair FOXP3’s DNA-binding ability (fig. S8A and B). Together, these findings suggest that the regulatory role of CXXC1 in T_reg_ cells is mediated through its effect on H3K4me3 deposition rather than altering FOXP3’s binding to DNA.

## Discussion

T_reg_ cells specifically express the transcription factor FOXP3, which is essential for maintaining T_reg_ lineage stability and suppressive function^53^. However, FOXP3 alone is insufficient to fully regulate the transcriptional signature and functionality of T_reg_ cells; its interaction with protein partners is crucial for this regulation. In this study, we identify CXXC1 as a critical epigenetic regulator and functional cofactor of FOXP3. Although CXXC1 is not required for FOXP3’s DNA-binding activity, as evidenced by similar FOXP3 binding patterns in WT and *Cxxc1*-deficient T_reg_ cells (CUT&Tag analysis), it plays an essential role in maintaining H3K4me3 modifications at FOXP3 target loci. By acting as both an epigenetic regulator and a FOXP3 cofactor, CXXC1 ensures the stability of the T_reg_ transcriptional program, highlighting its pivotal role in preserving T_reg_ cell functionality and immune homeostasis.

FOXP3 is known to promote histone H3 acetylation at the promoters and enhancers of its target genes, such as *Il2ra*, *Ctla4*, and *Tnfrsf18*, following T_reg_ cell activation, thereby functioning as a transcriptional activator^54,55^. Conversely, FOXP3 can act as a repressor by silencing target genes like *Il2* and *Ifng* through the induction of histone H3 deacetylation, mediated by the recruitment of histone deacetylases and transcriptional co-repressors^10,56^. Additionally, FOXP3 exerts this repression by recruiting the Ezh2-containing polycomb repressive complex to target genes during activation, as FOXP3-repressed genes are associated with H3K27me3 deposition and reduced chromatin accessibility^11,57^. Recent studies have begun to explore the biological significance of H3K4me3 breadth, revealing a positive correlation between H3K4me3 breadth and gene expression ^50,58,59^. These studies also suggest that H3K4me3 breadth contributes to defining specific cell identities during development and disease, including systemic autoimmune diseases like systemic lupus erythematosus and various cancers ^48,60–63^. Our study demonstrates that the loss of *Cxxc1* leads to reduced H3K4me3 levels, predominantly in genes with broader peaks, such as *Il2ra, Tnfrsf18, Ctla4 and Icos*, directly impairing their suppressive function in T_reg_ cells. Although our research primarily focused on the role of CXXC1 in T_reg_ cells, it is plausible that similar mechanisms may be operative in other cell types.

The T_reg_-specific deletion of *Cxxc1* leads to a rapid and fatal autoimmune disorder, characterized by systemic inflammation and tissue damage, underscoring the essential role of CXXC1 in maintaining immune self-tolerance within T_reg_ cells. Interestingly, despite this severe phenotype, our findings show that H3K4me1 levels were comparable between WT and *Cxxc1*-deficient T_reg_ cells. In contrast, MLL4 is critical for T_reg_ cell development in the thymus, primarily by regulating H3K4me1, though it is not required for peripheral T_reg_ cell function^15^. Numerous studies investigating the effectors of T_reg_ cell-mediated suppression have identified a wide range of molecules and mechanisms. These include the upregulation of the inhibitory co-stimulatory receptor CTLA-4, which initiates inhibitory signaling; the sequestration of the T-cell growth factor IL-2 via CD25; the secretion of inhibitory cytokines, such as interleukin (IL)-10, IL-35 and TGF-β and the activity of ectoenzymes CD39 and CD73 on the T_reg_ cells surface, which convert extracellular ATP, a potent pro-inflammatory mediator, into its anti-inflammatory counterpart, adenosine^42,64^. Mechanistically, we demonstrated that CXXC1 interacts with FOXP3 to regulate T_reg_ cell function by trimethylating H3K4 at broad H3K4me3 domains of multiple genes involved in suppressive functions such as *Il2ra, Nt5e, and Ctla4*. Additionally, MLL1, another KMT, controls T_reg_ cell activation and function by specifically regulating H3K4 trimethylation at genes encoding key T_reg_-related molecules such as *Tigit*, *Klrg1*, *Tbx21*, *Cxcr3*, and serves as a crucial epigenetic regulator in establishing a stable Th1-T_reg_ lineage^14^.

Our findings demonstrate that *Cxxc1*-deficient T_reg_ cells exhibit reduced H3K4me3 levels at T_reg_-specific loci, indicating that CXXC1 plays a critical role in regulating this epigenetic modification. This is consistent with previous studies showing that CXXC1 acts as a non-catalytic component of the Set1/COMPASS complex ^65–68^, which includes the H3K4 methyltransferases SETD1A and SETD1B, the primary enzymes responsible for H3K4 trimethylation. Given this, we propose that CXXC1 supports H3K4me3 deposition in T_reg_ cells by interacting with and stabilizing the activity of the Set1/COMPASS complex. Further studies are required to directly investigate the interactions between CXXC1 and these methyltransferases in the T_reg_ cell context.

Our results offer provide novel insights into the suppressive functions, heterogeneity, and regulatory mechanisms of T_reg_ cells. Maintaining T_reg_ cell homeostasis and function remains a challenge in harnessing T_reg_ cells for the treatment of autoimmune diseases and the prevention of graft rejection.

## Competing interests

The authors declare that they have no competing interests.

## Acknowledgments

We thank B. Li (Shanghai Jiao Tong University School of Medicine, Shanghai, China) for providing *Foxp3*^YFP-Cre^ mice, X.Y. Zhou(Chinese Academy of Science (CAS), Beijing, China) for providing the FOXP3 CUT&Tag antibody, and X. L. Liu (Shanghai Institutes for Biological Sciences, Chinese Academy of Sciences) for his generous gifts of cell lines. We thank Y. Huang, Y. Li, J. Wan, C.Guo, and Z. Lin from the Core Facilities, Zhejiang University School of Medicine for their technical support and S. Hong, Y. Ding, H. Jin, Q. Wang, and X. Zhang from Animal Facilities, Zhejiang University, for feeding the mice.

## Footnotes

This work was supported by grants from the National Natural Science Foundation of China (32341002, 32030035, 32321002, 32100693, and 32270839), the National Key R & D Program of China (2023YFA1800202, 2024YFF0728703, 2022YFA1103702, 2022YFA1103200), the Zhejiang Provincial Natural Science Foundation of China (LZ21C080001, LZ23C070003), Science and Technology Innovation 2030-Major Project (2021ZD0200405), Key project of the Experimental Technology Program of Zhejiang University (SZD202203), and the Fundamental Research Funds for the Central Universities (226-2024-00161).

## Author contributions

L.W. and L.S.: supervision, conceptualization, project administration, and writing—review and editing. L.W., L.S. and X.M: funding acquisition. X.M: investigation, methodology, project administration, and writing— original draft. Y.Z.and K.L.: bioinformatics analysis. Y. W., X.L., J.C., C.L., Y.Z., S.W., S.T., Q.X., L.D., X.S., and X.G.: investigation and methodology.

---

**Supplementary Figure 1.**
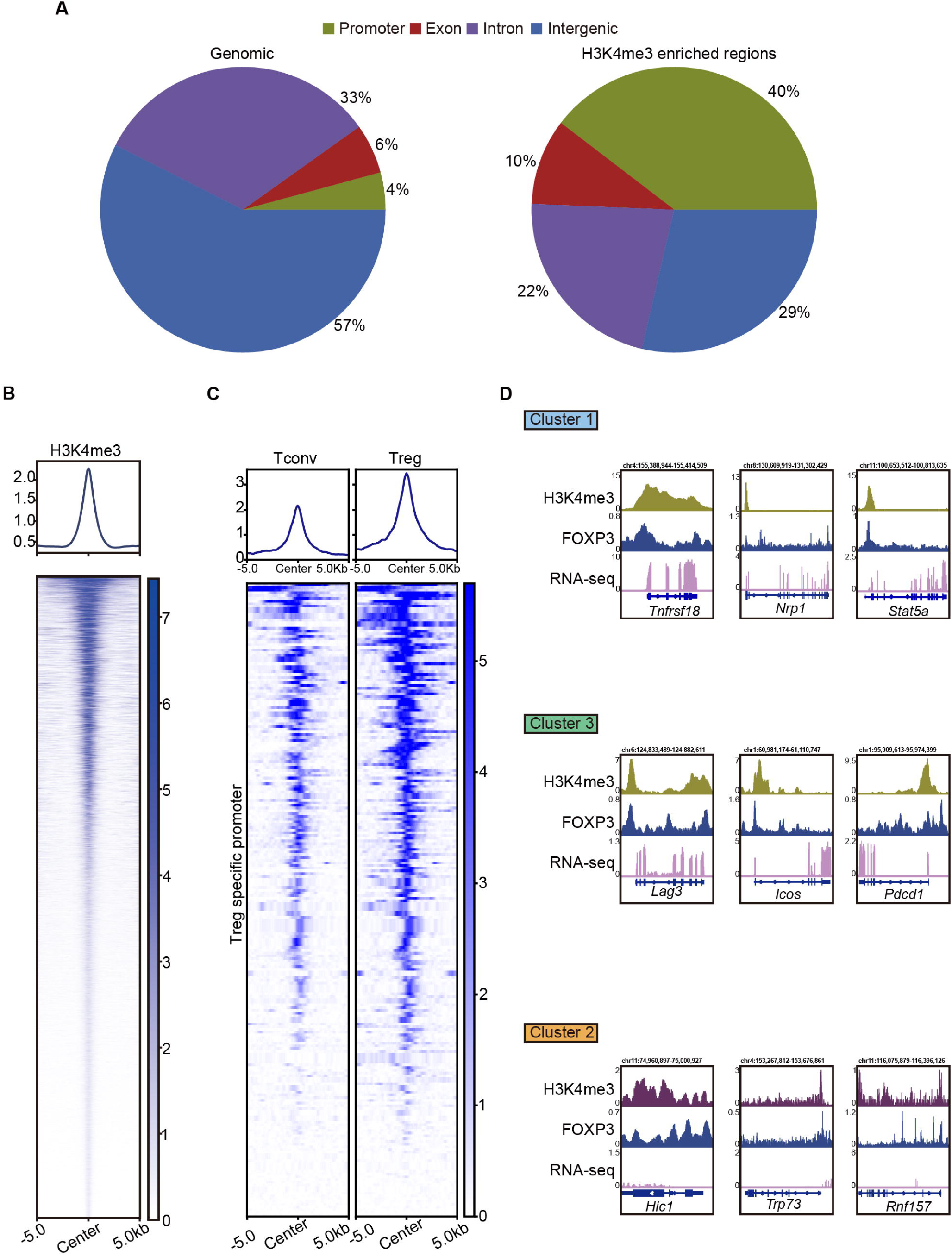

**Supplementary Figure 2.**
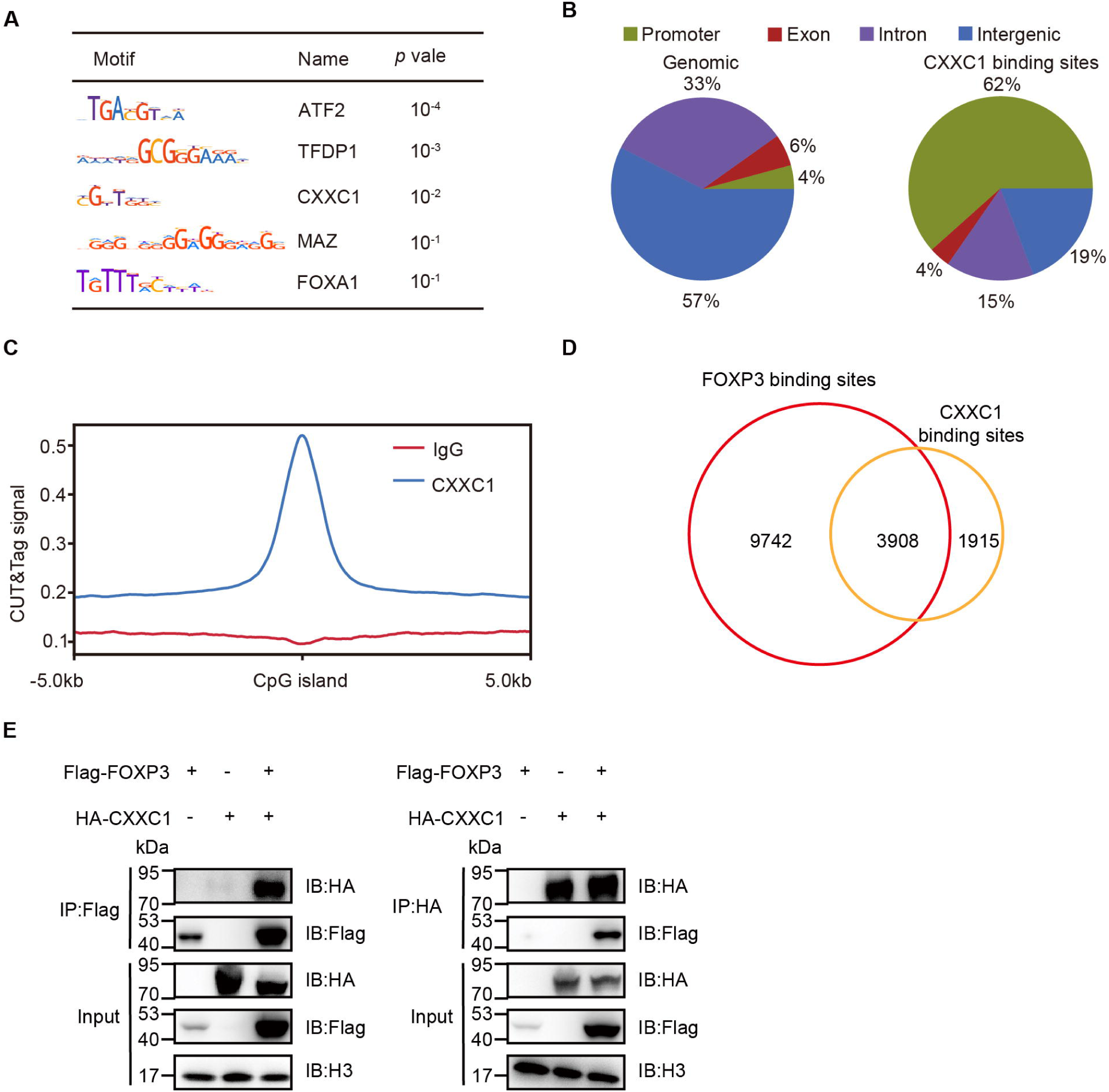

**Supplementary Figure 3.**
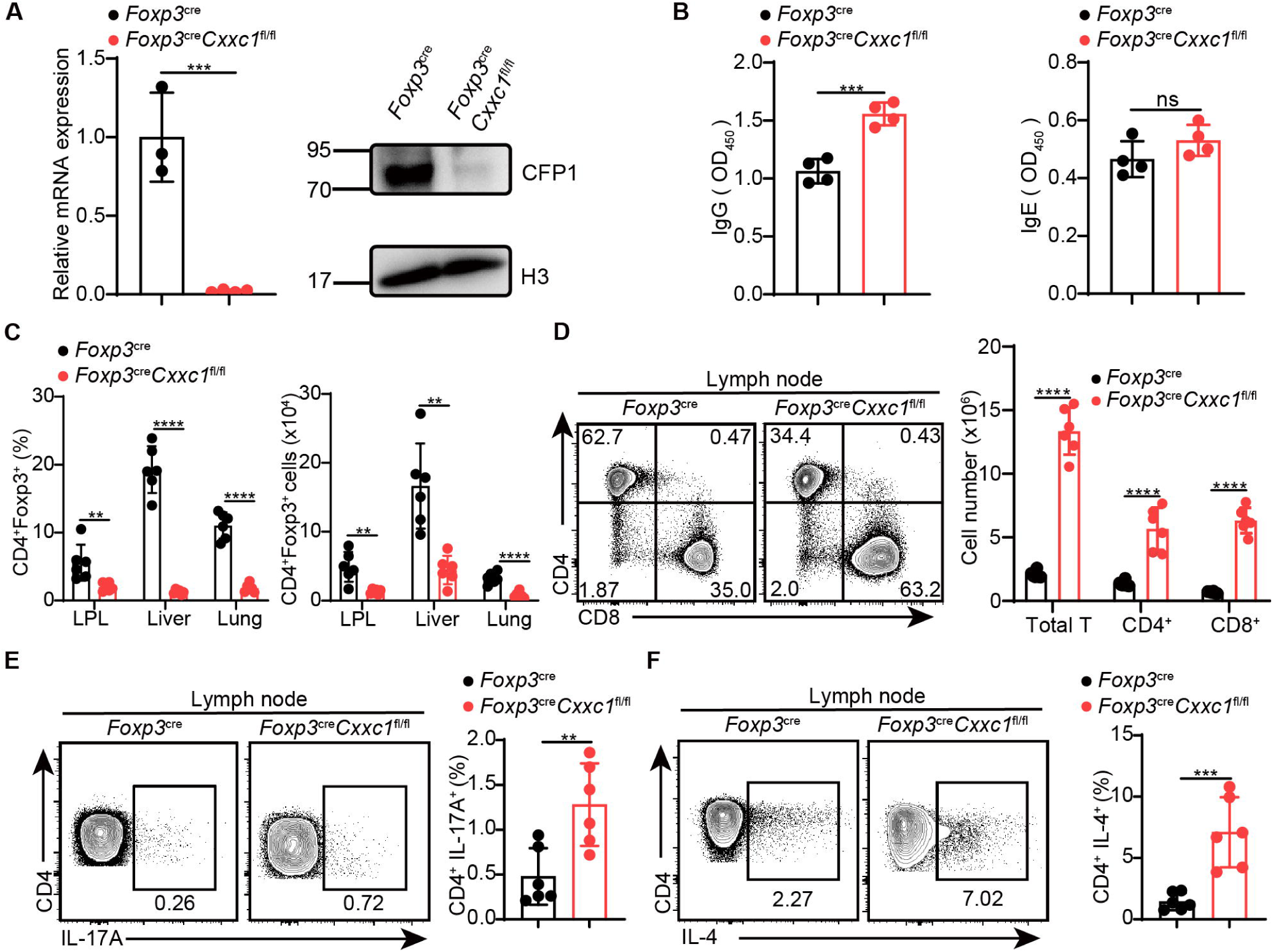

**Supplementary Figure 4.**
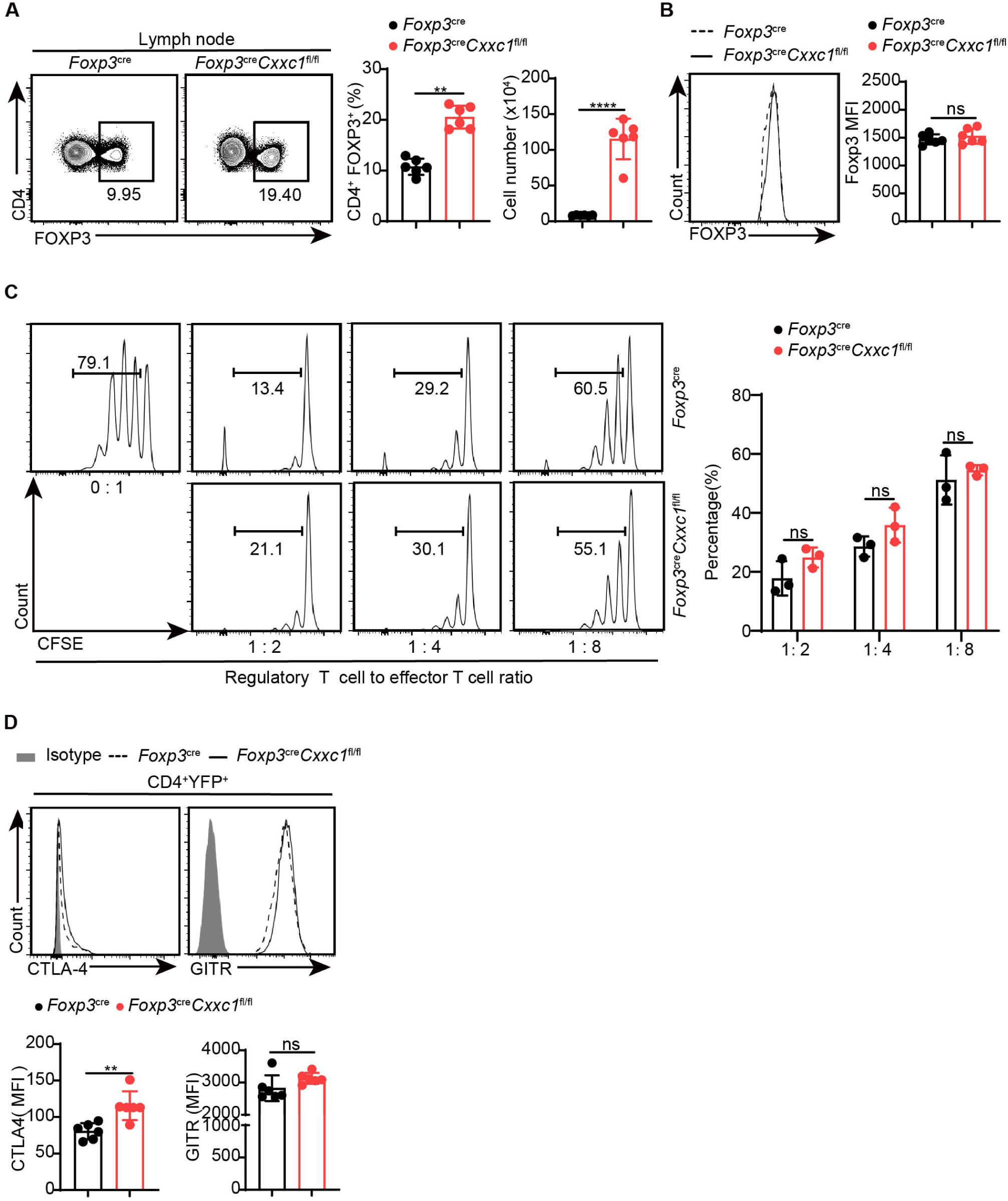

**Supplementary Figure 5.**
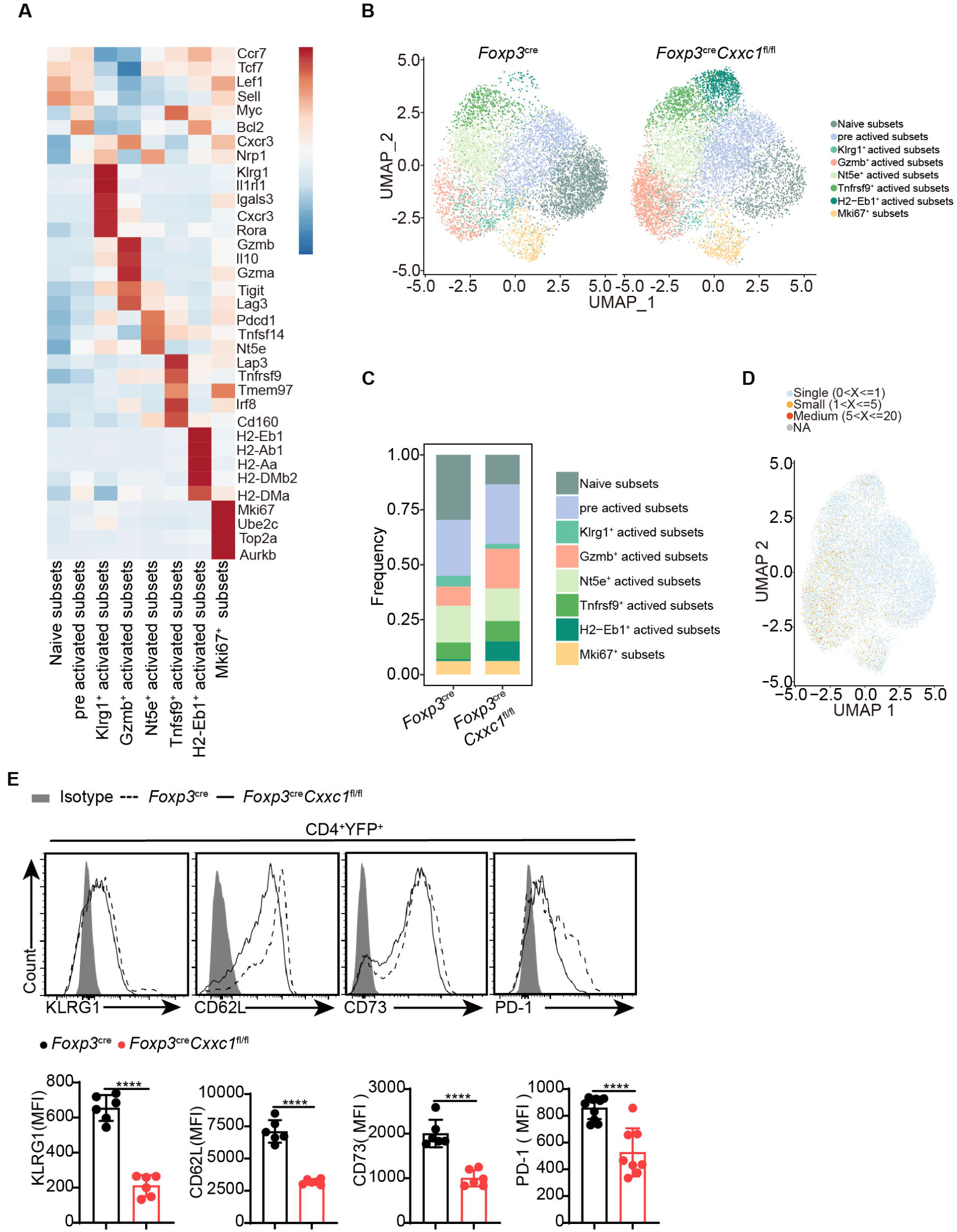

**Supplementary Figure 6.**
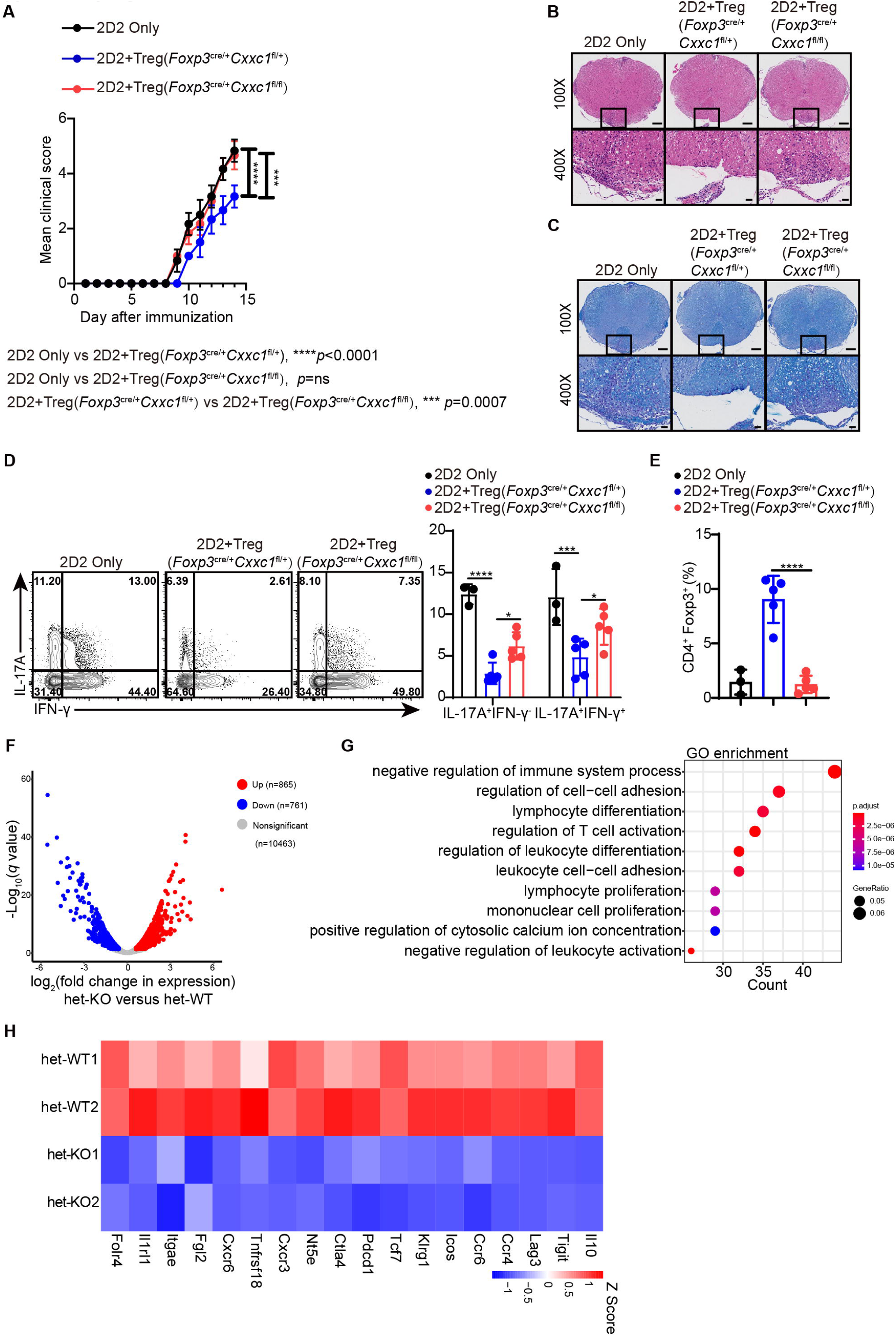

**Supplementary Figure 7.**
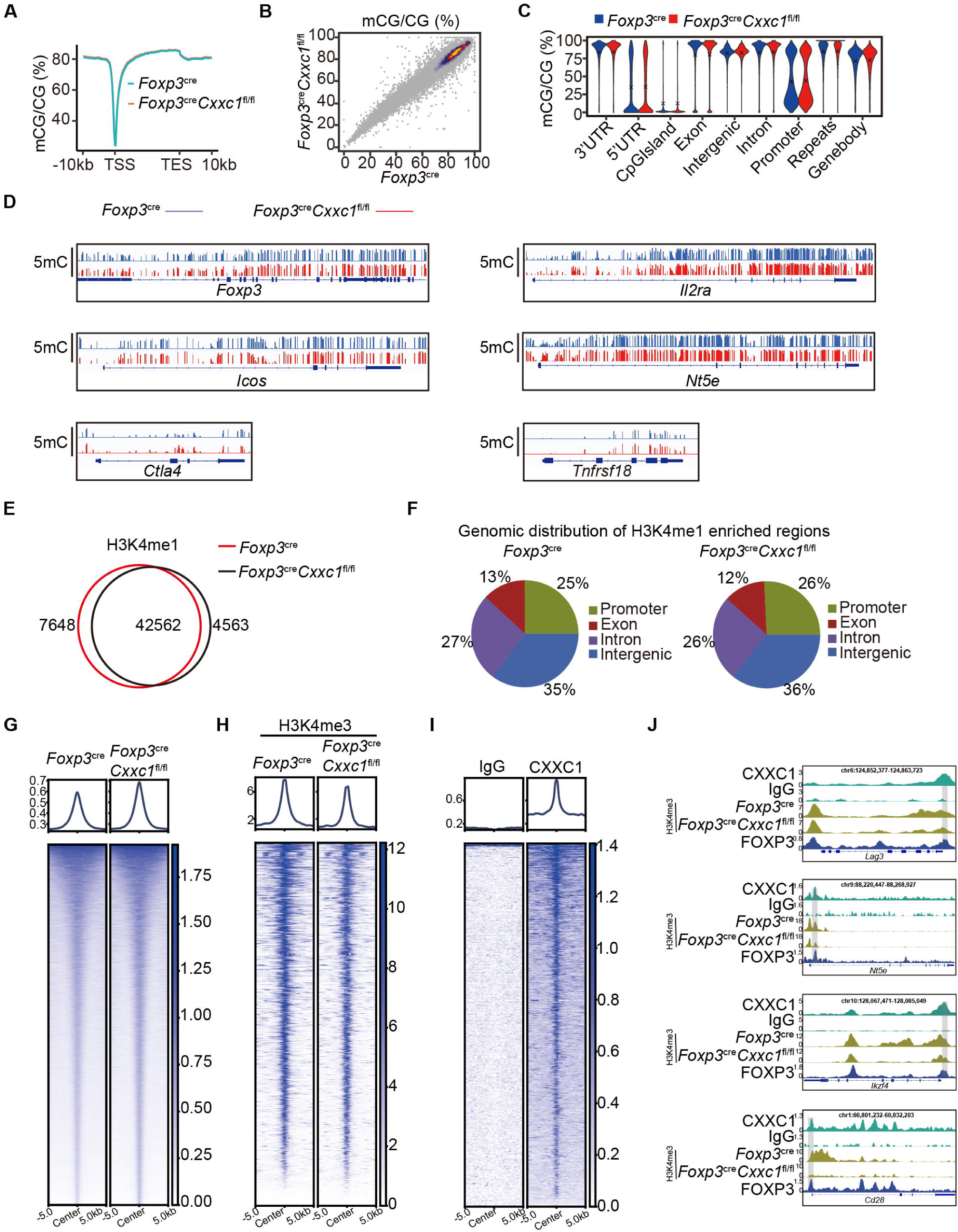

**Supplementary Figure 8.**
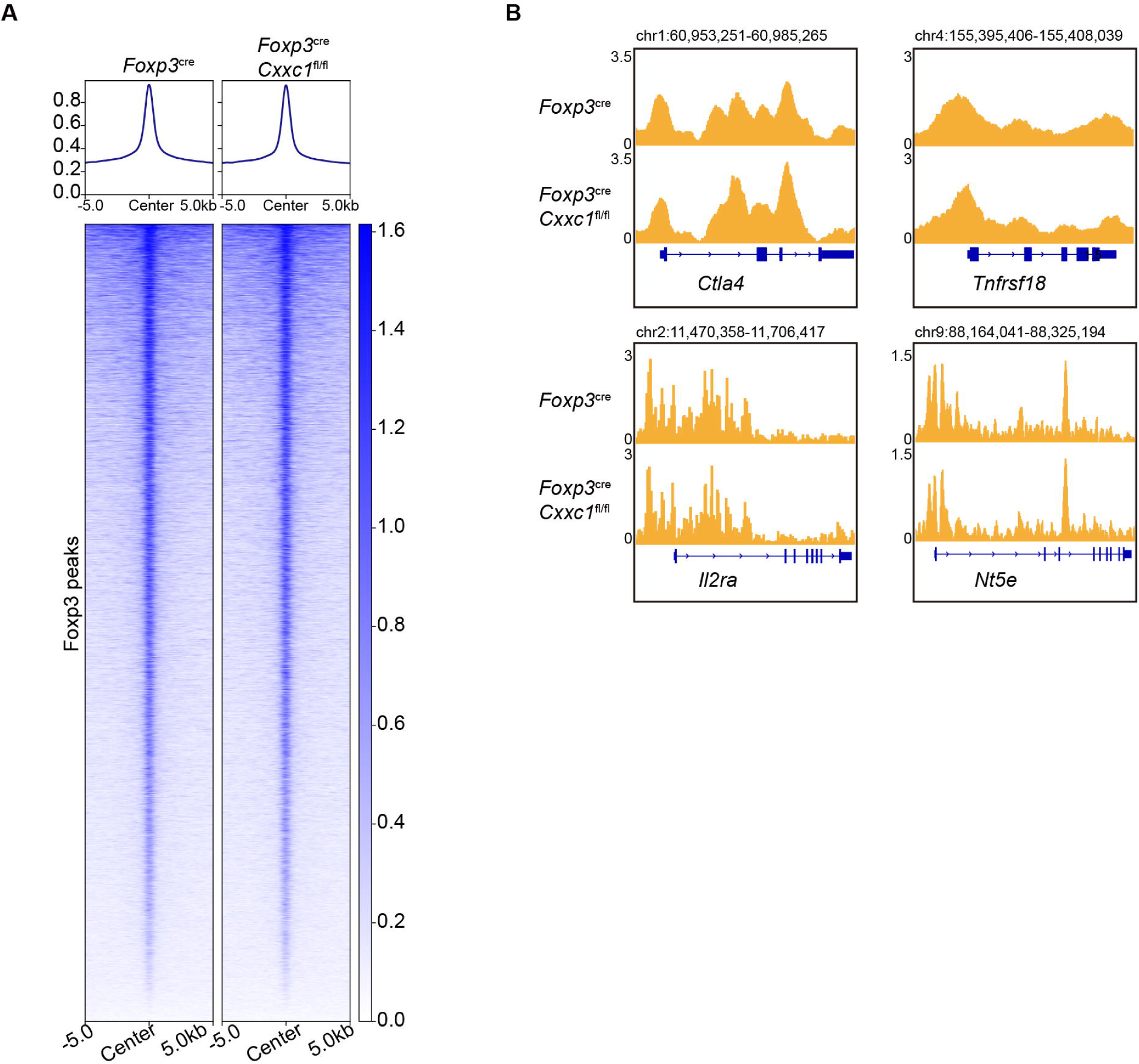

## Notes

### Competing Interest Statement

The authors have declared no competing interest.

### Summary of Updates

Figure 1 revised;Figure S1 revised;Figure 3 revised;Figure S3 revised;Figure 4 revised;Figure 6 revised;Figure S6 revised;Figure S8 revised;the text corresponding to the following figures has also been revised: Figure 1, Figure S1, Figure 3, Figure S3, Figure 4, Figure 6, Figure S6, and Figure S8. Supplemental Table updated.

