## Supplemental information for "CXXC-finger protein 1 associates with FOXP3 to stabilize homeostasis and suppressive functions of regulatory T cells"

**Figure S1. H3K4me3 is required for FOXP3-dependent gene activation in T_reg_ cells.**

(**A**)Genomic distribution of random and H3k4me3 enriched peaks in T_reg_ cells.

(**B**)Heatmap showing enrichment of H3K4me3 surrounding the TSS. The top panel of the profile plots shows the average CUT&Tag signals.

(**C**) Heatmap showing H3K4me3 enrichment at Treg-specific gene loci in Tconv and T_reg_ cells. The top panel of the profile plots shows the average CUT&Tag signals.

(**D**)Representative Genome Browser view showing RNA-seq, FOXP3 binding, and H3K4-me3/H3K27me3 enrichment in T_reg_ cells.

**Figure S2. FOXP3 interacts with CXXC1.**

(**A**) Motif enrichment analysis of overlap peaks in H3K4me3 enriched regions and FOXP3 binding sites.

(**B**)Genomic distribution of control and CXXC1-binding sites in T_reg_ cells.

(**C**)Average CXXC1 CUT&Tag signals around CpG island in T_reg_ cells. IgG was used as the control.

(**D**)Venn diagrams showing the overlap of FOXP3 binding sites and CXXC1 binding sites in T_reg_ cells.

(**E**)Physical interaction between CXXC1 and FOXP3 in HEK 293T cells transiently transfected with expression constructs. IP, immunoprecipitation.

**Figure S3. Disrupted immune homeostasis in *Foxp3*^cre^*Cxxc1*^fl/fl^ mice.**

**(A)**Analysis of *Cxxc1* mRNA (left) and protein (right) expression in CD4^+^YFP^+^ T_reg_ cells from *Foxp3*^cre^ and *Foxp3*^cre^*Cxxc1*^fl/fl^ mice.

**(B)**Quantification of serum IgG and IgE from *Foxp3*^cre^ and *Foxp3*^cre^*Cxxc1*^fl/fl^ mice(n=4).

**(C)**Quantification of the frequencies and absolute numbers of CD4⁺ Foxp3⁺ T_reg_ cells in the small intestinal lamina propria (LPL), liver, and lung of *Foxp3*^cre^ and *Foxp3*^cre^*Cxxc1*^fl/fl^ mice (n=6).

(**D**)Flow cytometry analysis of CD4 and CD8 expression in peripheral lymph nodes from *Foxp3*^cre^ and *Foxp3*^cre^*Cxxc1*^fl/fl^ mice. Absolute cell counts of Total T, CD4^+,^ and CD8^+^ T-cell subsets are shown (n=6).

(**E**, **F)**Lymph node cells from *Foxp3*^cre^ and *Foxp3*^cre^*Cxxc1*^fl/fl^ mice were stimulated ex vivowith PMA + ionomycin for 4 hours and analyzed for IL-17A^+^ and IL-4^+^ CD4^+^YFP^-^ T ce-lls by flow cytometry. Percentages of IL-17A^+^ (n=4) and IL-4^+^ (n=6) CD4^+^ T cells in ly-mphnodes from *Foxp3*^cre^ and *Foxp3*^cre^*Cxxc1*^fl/fl^ mice.

Error bars show mean ± SD. *P* values are determined by a two-tailed Student’s *t*-test (A-D). (***P*<0.01, ****P* < 0.001, *****P*<0.0001). The flow cytometry and western blot results represent at least three independent experiments.

**Figure S4. The number of T_reg_ cells deficient in CXXC1 did not decrease.**

(**A**)Flow cytometry analysis of CD4^+^FOXP3^+^ T_reg_ cells in lymph nodes of *Foxp3*^cre^ and *F-oxp3*^cre^*Cxxc1*^fl/fl^ mice. On the right, the proportion and number of T_reg_ cells are shown (n=6).

(**B**)Representative figure showing FOXP3 protein expression in CD4^+^ T cells from the lymph nodes (n=10).

(**C**)Suppression of CFSE-labelled Tn cell proliferation by different ratios of CD4^+^YFP^+^ T_reg_ cells from *Foxp3*^cre^ and *Foxp3*^cre^*Cxxc1*^fl/fl^ mice. On the right, the percentage of proliferated responding T cells is presented (n=3).

(**D**)Expression of T_reg_ signature molecules in *Foxp3*^cre^ and *Foxp3*^cre^*Cxxc1*^fl/fl^ mice (n=4 CCR7, n=6 CTLA-4, n=6 GITR).

Error bars show mean ± SD. *P* values are determined by a two-tailed Student’s *t*-test (A-D). (ns, not significant. **P*<0.05, ***P*<0.01, ****P* < 0.001, *****P*<0.0001). The flow cytometry results are representative of three independent experiments.

**Figure S5. Single-cell transcriptomics reveals distinct Treg cell populations.**

(**A**)Heatmap displaying the Z scores for the average expression of T_reg_-specific genes across each cluster.

(**B**)UMAP plot showing clusters identified by variable gene expression in sorted CD4^+^YFP^+^ T_reg_ cells from *Foxp3*^cre^ and *Foxp3*^cre^*Cxxc1*^fl/fl^ mice.

(**C**)Relative proportions of T_reg_ cell subpopulations in *Foxp3*^cre^ and *Foxp3*^cre^*Cxxc1*^fl/fl^, revealing heterogeneity.

(**D**)Density and clonotype richness across T_reg_ cell clusters, with colors indicating clone size.

(**E**)Representative flow plots and quantified expression of KLRG1, CD62L, CD73, and PD-1 in CD4^+^YFP^+^ T_reg_ cells from *Foxp3*^cre^ and *Foxp3*^cre^*Cxxc1*^fl/fl^ mice (n=6 KLRG1, n=6 CD62L, n=6 CD73, n=8 PD-1).

Error bars show mean ± SD. *P* values are determined by a two-tailed Student’s *t*-test (E). (*****P*<0.0001).

**Figure S6. *Cxxc1* deficiency impairs T_reg_ Cell suppressive function and aggravates EAE severity.**

(**A**)Mean clinical scores for EAE in *Rag1^−/−^* recipients of 2D2 CD4^+^ T cells, either alone or in combination with *Foxp3*^Cre/+^ *Cxxc1*^fl/+^ or *Foxp3*^Cre/+^ *Cxxc1*^fl/fl^ female mice after immunization with MOG35–55, complete Freund’s adjuvant (CFA), and pertussis toxin (n=6).

(**B** and **C**)Representative histology of the spinal cord of *Rag1^−/−^* mice after EAE induction. Hematoxylin and eosin(H&E) staining (upper), Luxol fast blue (F&B) staining (lower). Scale bars, 50 μm (400×) and 200 μm (100×).

(**D**)Representative flow cytometry plots and quantification of the the percentages of IFNγ^+^ or IL-17A^+^ CD4^+^Vβ11^+^ T cells.

(**E**)Statistical analysis of the percentage CD4^+^ FOXP3^+^ T_reg_ cell in CNS tissues 14 days after EAE induction.

(**F**)Volcano plot showing the expression of genes in CD4^+^YFP^+^ Treg cells from het-KO versus het-WT. Differential Expressed Genes (DEGs) whose expressions are 1.5-fold up-regulated or down-regulated, respectively.

(**G**)Gene set enrichment analysis identified downregulated hallmark pathways in CD4^+^YFP^+^ T_reg_ cells.

(**H**)Heatmap showing the relative accumulation of T_reg_ cell signature genes implicated in suppressive function; relative expression of genes (Z score) is displayed and color coded according to the score.

Error bars show mean ± SD. *P* values are determined by a two-tailed Student’s *t*-test (A, D and E). (**P*<0.05, ****P* < 0.001, *****P*<0.0001).

**Figure S7. The epigenetic program of T_reg_ cells.**

(**A**)Average CpG methylation in CD4^+^YFP^+^ cells from three-week-old *Foxp3*^cre^ and *Foxp3*^c-re^*Cxxc1*^fl/fl^ surrounding the transcription start site (TSS) and transcriptional terminal site (TES).

(**B**)Scatter plot showing methylation levels at 5 kb bins in wild-type (WT) and *Cxxc1*-deficient T_reg_ cells.

(**C**)A violin plot showing the methylation levels at different genomic regions in WT and *Cxxc1*-deficient T_reg_ cells.

(**D**)Genome browser view showing DNA methylation levels in WT and *Cxxc1*-null T_reg_ cells.

(**E**)Venn diagrams showing the overlap of H3K4me1 peaks in WT and *Cxxc1*-null T_reg_ cells.

(**F**)Genomic distribution of H3K4me1 peaks in WT and *Cxxc1*-null T_reg_ cells.

(**G**)Heatmap showing H3K4me1 enrichment in WT and *Cxxc1*-null T_reg_ cells. The top panel of the profile plots shows the average CUT&Tag signals.

(**H**-**I**)Heatmaps showing H3K4me3(H)and CXXC1(I) signals centered on broad domains. The top panel shows the average CUT&Tag signals around indicated domains.

(**J**)Representative genome browser view showing the enrichments of FOXP3, CXXC1, and H3K4me3 in T_reg_ cells.

**Figure S8. FOXP3 binding profile and enrichment in WT and *Cxxc1*-Null T_reg_ cells.**

(**A**)Heatmap showing FOXP3 CUT&Tag signals in WT and *Cxxc1*-null T_reg_ cells. The top panel of the profile plots shows the average CUT&Tag signals.

(**B**)Representative genome browser views showing enrichment of FOXP3 in WT and *Cxxc1*-null T_reg_ cells.

Supplementary Table S1. Quality control of CUT&Tag data.

| Sample | Total reads | Mapping efficiency | Correlation |
| --- | --- | --- | --- |
| WT-IgG | 28,433,877 | 97.42% |  |
| WT-CXXC1-rep1 | 37,605,589 | 98.00% | 0.96 |
| WT-CXXC1-rep2 | 40,625,734 | 97.64% |  |
| WT-H3K4me1-rep1 | 28,324,337 | 98.61% | 0.99 |
| WT-H3K4me1-rep2 | 23,814,172 | 99.20% |  |
| cKO-H3K4me1-rep1 | 26,325,592 | 98.74% | 0.97 |
| cKO-H3K4me1-rep2 | 22,378,739 | 99.20% |  |
| WT-H3K4me3-rep1 | 14,464,350 | 97.01% | 0.92 |
| WT-H3K4me3-rep2 | 41,233,666 | 99.06% |  |
| cKO-H3K4me3-rep1 | 10,423,744 | 97.86% | 0.88 |
| cKO-H3K4me3-rep2 | 22,365,710 | 98.88% |  |

Supplementary Table S2. Quality control of WGBS data.

| Sample | Total reads | Mapping efficiency | CpG methylation level |
| --- | --- | --- | --- |
| WT -WGBS | 461,596,431 | 77.0% | 79.4% |
| cKO -WGBS | 459,215,408 | 72.8% | 80.3% |
